# BBX proteins promote HY5-mediated UVR8 signaling in Arabidopsis

**DOI:** 10.1101/2021.10.14.464399

**Authors:** Roman Podolec, Timothée B. Wagnon, Manuela Leonardelli, Henrik Johansson, Roman Ulm

## Abstract

Plants undergo photomorphogenic development in the presence of light. Photomorphogenesis is repressed by the E3 ubiquitin ligase CONSTITUTIVELY PHOTOMORPHOGENIC1 (COP1), which binds substrates through their valine-proline (VP) motifs. The UV RESISTANCE LOCUS8 (UVR8) photoreceptor senses UV-B and inhibits COP1 through cooperative binding of its own VP motif mimicry and its photosensing core to COP1, thereby preventing COP1 binding to substrates, including the bZIP transcriptional regulator ELONGATED HYPOCOTYL5 (HY5). As a key promoter of visible light and UV-B photomorphogenesis, HY5 functions together with the B-box family transcription factors BBX20–22 that were recently described as HY5 rate-limiting coactivators under red light. Here we describe a hypermorphic *bbx21-3D* mutant with enhanced photomorphogenesis, which carries a proline-314 to leucine mutation in the VP motif that impairs interaction with and regulation through COP1. We show that BBX21 and BBX22 are UVR8-dependently stabilized after UV-B exposure, which is counteracted by a repressor induced by HY5/BBX activity. *bbx20 bbx21 bbx22* mutants under UV-B are impaired in hypocotyl growth inhibition, photoprotective pigment accumulation, and expression of several HY5-dependent genes. We conclude that BBX20–22 importantly contribute to HY5 activity in a subset of UV-B responses, but that additional, presently unknown coactivators for HY5 are functional in early UVR8 signaling.

## INTRODUCTION

Plants integrate light signals through a number of photoreceptor-initiated signaling pathways (Kami et al., 2010; Galvao and Fankhauser, 2015; Demarsy et al., 2018; Podolec et al., 2021a), which enables optimized growth in changing environments. When exposed to light, including ultraviolet-B (UV-B), seedlings undergo a developmental program termed photomorphogenesis, characterized by features such as hypocotyl growth inhibition, anthocyanin and flavonoid pigment accumulation, and cotyledon expansion (Kami et al., 2010; Galvao and Fankhauser, 2015). In Arabidopsis (*Arabidopsis thaliana*), UV-B radiation (280–315 nm) is sensed by the photoreceptor UV RESISTANCE LOCUS8 (UVR8) that induces a signaling pathway resulting in UV-B acclimation and thus enhanced UV-B tolerance (Kliebenstein et al., 2002; Favory et al., 2009; Rizzini et al., 2011; Podolec et al., 2021a; Rai et al., 2021). UVR8 is homodimeric in its inactive ground state, but monomerizes and becomes activated upon UV-B photon reception (Rizzini et al., 2011). A valine-proline (VP) motif in activated UVR8 represses the activity of the E3 ubiquitin ligase CONSTITUTIVELY PHOTOMORPHOGENIC1 (COP1) by competitively interacting with the binding site of COP1 substrates (Favory et al., 2009; Cloix et al., 2012; Yin et al., 2015; Lau et al., 2019).

Prominent among COP1 substrates is ELONGATED HYPOCOTYL5 (HY5), a key photomorphogenesis-promoting bZIP transcriptional regulator that induces expression of many UV-B-activated genes (Ulm et al., 2004; Brown et al., 2005; Oravecz et al., 2006). Upon UV-B exposure, *HY5* is transcriptionally induced and the HY5 protein is stabilized (Ulm et al., 2004; Brown et al., 2005; Favory et al., 2009; Huang et al., 2013). However, the HY5 mode of action in UV-B signaling is not fully understood as it lacks a functional transcriptional activation domain (Ang et al., 1998; Stracke et al., 2010; Burko et al., 2020). Nevertheless, HY5 binds the promoters of many UV-B-induced genes and is required for their UV-B-induced expression (Ulm et al., 2004; Brown et al., 2005; Oravecz et al., 2006; Brown and Jenkins, 2008; Stracke et al., 2010; Binkert et al., 2014). Thereby, HY5 promotes UV-B responses such as hypocotyl growth inhibition, anthocyanin and flavonoid pigment accumulation, and ultimately stress acclimation (Brown et al., 2005; Oravecz et al., 2006; Favory et al., 2009; Huang et al., 2012). HY5-regulated genes include those encoding REPRESSOR OF UV-B PHOTOMORPHOGENESIS1 (RUP1) and RUP2, which are crucial negative feedback regulators facilitating UVR8 ground state reversion by redimerization (Gruber et al., 2010; Heijde and Ulm, 2013). In agreement, *rup1 rup2* display strong UV-B hypersensitivity, further supported by the phenotype of the UVR8^G101S^ mutant variant that is constitutively monomeric in vivo and underlies the enhanced UV-B photomorphogenic phenotype of the *uvr8-17D* mutant allele (Podolec et al., 2021b).

Recently, the transactivation domain-containing, B-box zinc-finger transcription factors BBX20–22 were characterized as coregulators of HY5 under red light, interacting with HY5 and providing it with transcriptional activity (Bursch et al., 2020). BBX20–22, as well as BBX23, another member of class IV of the B-box family, have been described as positive regulators of photomorphogenesis (Datta et al., 2007; Chang et al., 2008; Datta et al., 2008; Chang et al., 2011; Fan et al., 2012; Wei et al., 2016; Zhang et al., 2017), whereas two other members of the subfamily, namely BBX24 and BBX25, play a repressive role (Indorf et al., 2007; Yan et al., 2011; Jiang et al., 2012; Gangappa et al., 2013; Crocco et al., 2015). In darkness, BBX20–25 are degraded via COP1, whereas they are stabilized in the light (Indorf et al., 2007; Chang et al., 2011; Fan et al., 2012; Jiang et al., 2012; Gangappa et al., 2013; Xu et al., 2016; Zhang et al., 2017; Job et al., 2018). BBX21, BBX24, and BBX25 contain experimentally defined COP1-interacting VP motifs, similar to other COP1 targets (Holm et al., 2001; Yan et al., 2011; Lau et al., 2019; Bursch et al., 2020). Moreover, HY5 has been suggested to repress BBX22 accumulation in an unknown manner (Chang et al., 2011). Whereas the negative regulator BBX24 has been implicated in UV-B signaling by suppressing HY5 activity (Jiang et al., 2012), it is not known whether the class IV BBX family positive regulators play a role in UV-B signaling in Arabidopsis. However, the involvement of functional homologs of class IV BBX positive regulators in UV-B signaling and anthocyanin biosynthesis has been reported in apple (*Malus × domestica*) in transgenic overexpression and suppression lines (Bai et al., 2014; Fang et al., 2019).

In this work, we identified and characterized a novel gain-of-function mutant of BBX21, namely *bbx21-3D*, that contains a hypermorphic proline-314 to leucine (P314L) mutation in its VP motif. Phenotypic analyses suggested that the enhanced photomorphogenesis of this mutant is linked to increased activity of BBX21^P314L^ and impaired negative regulation by COP1. We further explored the role of the BBX20–22 subfamily in UV-B photomorphogenesis and found that BBX proteins are regulated at both transcriptional and post-translational levels under UV-B. We uncovered a HY5/BBX-mediated negative feedback mechanism affecting BBX protein stability. Moreover, a *bbx20 bbx21 bbx22* mutant showed a lack of flavonoid and anthocyanin accumulation, impaired inhibition of hypocotyl elongation, and reduced marker gene expression under UV-B. Collectively, our work reveals that BBX20–22 play an important role in promoting HY5-mediated UV-B responses.

## RESULTS

### *bbx21-3D* is a gain-of-function mutant of *BBX21* with an enhanced photomorphogenic response

In a hypocotyl length–based EMS mutant screen for altered UV-B photomorphogenesis (Podolec et al., 2021b), we discovered a mutant with an enhanced photomorphogenic response under white light with supplemental UV-B (mutant *bbx21-3D* in Fig. 1A,B). However, the manifestation of this phenotype under white light in the absence of supplemental UV-B indicated that it was not specific to UV-B (Fig. 1A,B). A causative transition mutation in *AT1G75540* (*BBX21*) was identified by whole-genome sequencing of a bulk segregant population; this mutation alters the VP motif of BBX21, namely proline-314 to leucine, resulting in a BBX21^P314L^ variant (Supplemental Fig. S1A). The novel mutant allele showed a dominant hypocotyl phenotype (Fig. 1C), and hence was named *bbx21-3D*. To verify that the enhanced photomorphogenic phenotype of *bbx21-3D* was caused by the BBX21^P314L^ mutation, we used CRISPR/Cas9 to knockout *BBX21* in the *bbx21-3D* background. Unlike *bbx21-3D*, a *BBX21* knockout allele in *bbx21-3D* showed an elongated hypocotyl phenotype comparable to the *bbx21-1* null allele as well as the *bbx21*^*KO*^ CRISPR/Cas9 knockout allele in the Col-0 wild-type background generated here (Supplemental Fig. S1B,C). These data support that BBX21^P314L^ is the causative mutation for the hypermorphic phenotype of *bbx21-3D*. To further confirm that the *bbx21-3D* phenotype is due to the loss of a functional VP motif, we used CRISPR/Cas9 to mutate the C-terminus of BBX21 in a wild-type background (Supplemental Fig. S1B). Indeed, a premature stop codon causing a C-terminal deletion removing the BBX21 VP motif mimicked the short-hypocotyl phenotype of *bbx21-3D* (Supplemental Fig. S1D), supporting that the VP motif is crucial to negatively regulate BBX21 activity.

**Figure 1.**
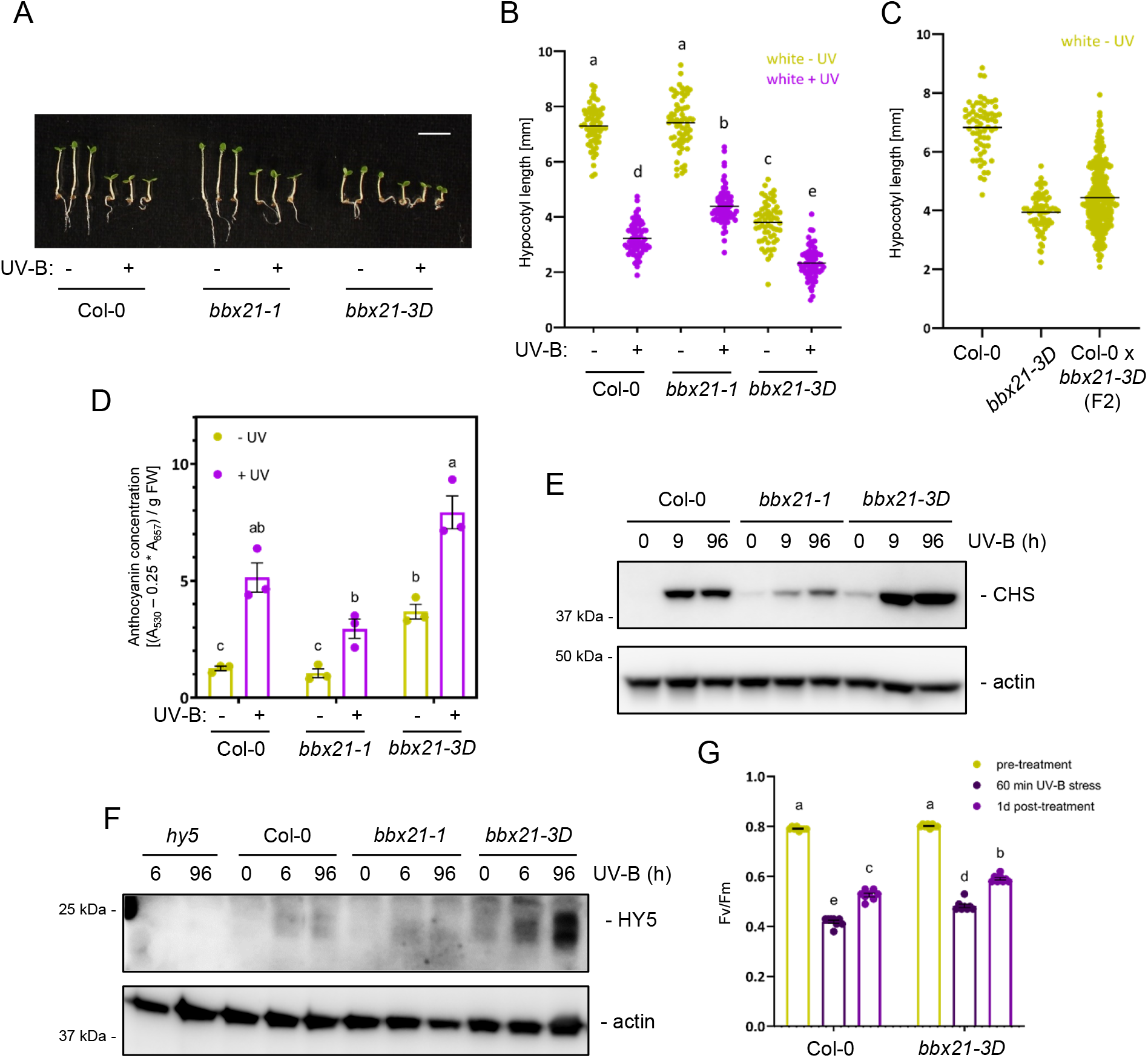
*bbx21-3D* is a hypermorphic allele of BBX21. A, Representative images of wild type (Col-0) and *bbx21-1* null and *bbx21-3D* gain-of-function mutant seedlings in white light supplemented with UV-B (+), or not (–). Scale bar indicates 5 mm. B, Quantification of hypocotyl length of seedlings as shown in (A). C, Quantification of hypocotyl lengths of Col-0, *bbx21-3D*, and segregating F2 seedlings of a Col-0 x *bbx21-3D* cross grown for 4 d in white light. D, Anthocyanin concentrations in seedlings as shown in (A). E, Immunoblot analysis of CHS and actin (loading control) levels in Col-0, *bbx21-1*, and *bbx21-3D* seedlings exposed to 9 h and 96 h of supplemental UV-B, or not (0). F, Immunoblot analysis of HY5 and actin (loading control) levels in Col-0, *bbx21-1, bbx21-3D* and *hy5* seedlings exposed to 6 h and 96 h of supplemental UV-B, or not (0). G, Maximum efficiency of photosystem II (Fv/Fm) in wild-type (Col-0) and *bbx21-3D* seedlings grown for 7d in white light (“pre-treatment”), exposed to 60 min of broadband UV-B (“60 min UV-B stress”), and left in white light for 1d of recovery (“1d post-treatment”). (B,C) Values of independent measurements and means as horizontal lines are shown; *n* > 60. (D) Values of independent measurements, means, and SEM are shown (*n* = 3). (G) Values of independent measurements, means, and SEM are shown (*n* = 8). Shared letters indicate no statistically significant difference between the means (P > 0.05).

Whereas *bbx21-3D* showed an enhanced photomorphogenic response in both –UV-B and +UV-B conditions, the *bbx21-1* null mutant exhibited slightly reduced hypocotyl growth inhibition in the presence of UV-B (Fig. 1A,B). Consistently, anthocyanin accumulation was enhanced in *bbx21-3D* in both –UV-B and +UV-B conditions compared to wild type, but was reduced relative to wild type under +UV-B in *bbx21-1* (Fig. 1D). UV-B-induced accumulation of CHALCONE SYNTHASE (CHS), a key enzyme for phenylpropanoid biosynthesis, was also enhanced in *bbx21-3D* relative to wild type, whereas it accumulated to lower levels in the *bbx21-1* null mutant (Fig. 1E). In *bbx21-3D*, UV-B-induced *HY5* activation was comparable to wild type, indicating that the *bbx21-3D* enhanced photomorphogenic phenotype was not associated with increased *HY5* transcript levels (Supplemental Fig. S2A). However, we observed an increase in HY5 protein accumulation in *bbx21-3D* compared to wild type in response to UV-B (Fig. 1F). BBX21^P314L^ thus likely promotes HY5 accumulation at the post-transcriptional level. It is also of note that *BBX21* mRNA levels were not elevated in *bbx21-3D* compared to wild type (Supplemental Fig. S2B). Moreover, whereas UV-B-induced expression of *RUP2* seemed unaffected by BBX21 (Supplemental Fig. S2C), several other UV-B marker genes such as *EARLY LIGHT-INDUCIBLE PROTEIN2* (*ELIP2*), *CHS*, and *FLAVANONE 3-HYDROXYLASE* (*F3H*) showed lower gene activation in *bbx21-1* compared to *bbx21-3D* (Supplemental Fig. S2D–F). Ultimately, *bbx21-3D* showed constitutive acclimation to UV-B, as its photosynthetic capacity was less affected by a UV-B stress compared to the wild type (Fig. 1G). Altogether, these data support a positive regulatory role of BBX21 in the UVR8 signaling pathway.

As BBX21 promotes photomorphogenesis under visible light (Datta et al., 2007; Xu et al., 2016), we tested the *bbx21-3D* mutant in darkness and red and blue light. In darkness, *bbx21-1* and *bbx21-3D* seedling phenotypes were comparable to wild type (Supplemental Fig. S3A,B). However, under both red and blue light, *bbx21-1* indeed showed a reduced photomorphogenic phenotype relative to wild type, whereas *bbx21-3D* showed an enhanced photomorphogenic phenotype, including hypocotyl growth inhibition and anthocyanin accumulation (Supplemental Fig. S3C–H). Overall, the contrasting phenotypes of *bbx21-3D* and *bbx21-1* support a positive regulatory role of BBX21 in photomorphogenic responses to both visible light and UV-B.

### BBX21^P314L^ function requires HY5 and shows increased activity in a both COP1-dependent and -independent manner

The VP motif of BBX21 is located close to the C-terminus of the protein (residues 313/314 of the 331 amino acid protein; Supplemental Fig. S1A,B). This motif assumedly mediates the interaction with COP1, leading to polyubiquitination and proteasomal degradation of BBX21 during skotomorphogenesis (Xu et al., 2016; Job et al., 2018; Lau et al., 2019; Bursch et al., 2020). In agreement, mutation of the VP motif to Ala-Ala (AA) increased the post-translational stability of BBX21 (Bursch et al., 2020), C-terminal deletion including the VP motif mimicked the *bbx21-3D* phenotype (Supplemental Fig. S1B,D), and the BBX21^P314L^ mutation resulted in hypermorphic phenotypes in all tested light conditions (Fig. 1 and Supplemental Fig. S3).

We thus tested the interactions of BBX21 and BBX21^P314L^ with HY5 and the C-terminal WD40-domain of COP1 (COP1^C340^) in yeast two-hybrid (Y2H) assays. Whereas interaction of BBX21 with COP1^C340^ was clearly detectable, BBX21^P314L^ interaction with COP1^C340^ was impaired (Fig. 2A). Interestingly, by contrast, the interaction between BBX21^P314L^ and HY5 was strongly enhanced compared to the BBX21–HY5 interaction (Fig. 2A). The intrinsic transcriptional activities (“auto-activation”) of BBX21^P314L^ and BBX21 fused to the LexA binding domain were comparable in the heterologous yeast system (Fig. 2B), making it unlikely that the *bbx21-3D* phenotype is due to intrinsically enhanced transcriptional activity of BBX21^P314L^. Nonetheless, the loss of interaction with COP1, and thus putative stabilization of BBX21^P314L^, and the enhanced BBX21^P314L^–HY5 interaction provide mutually non-exclusive mechanistic explanations for the *bbx21-3D* enhanced photomorphogenic phenotype.

**Figure 2.**
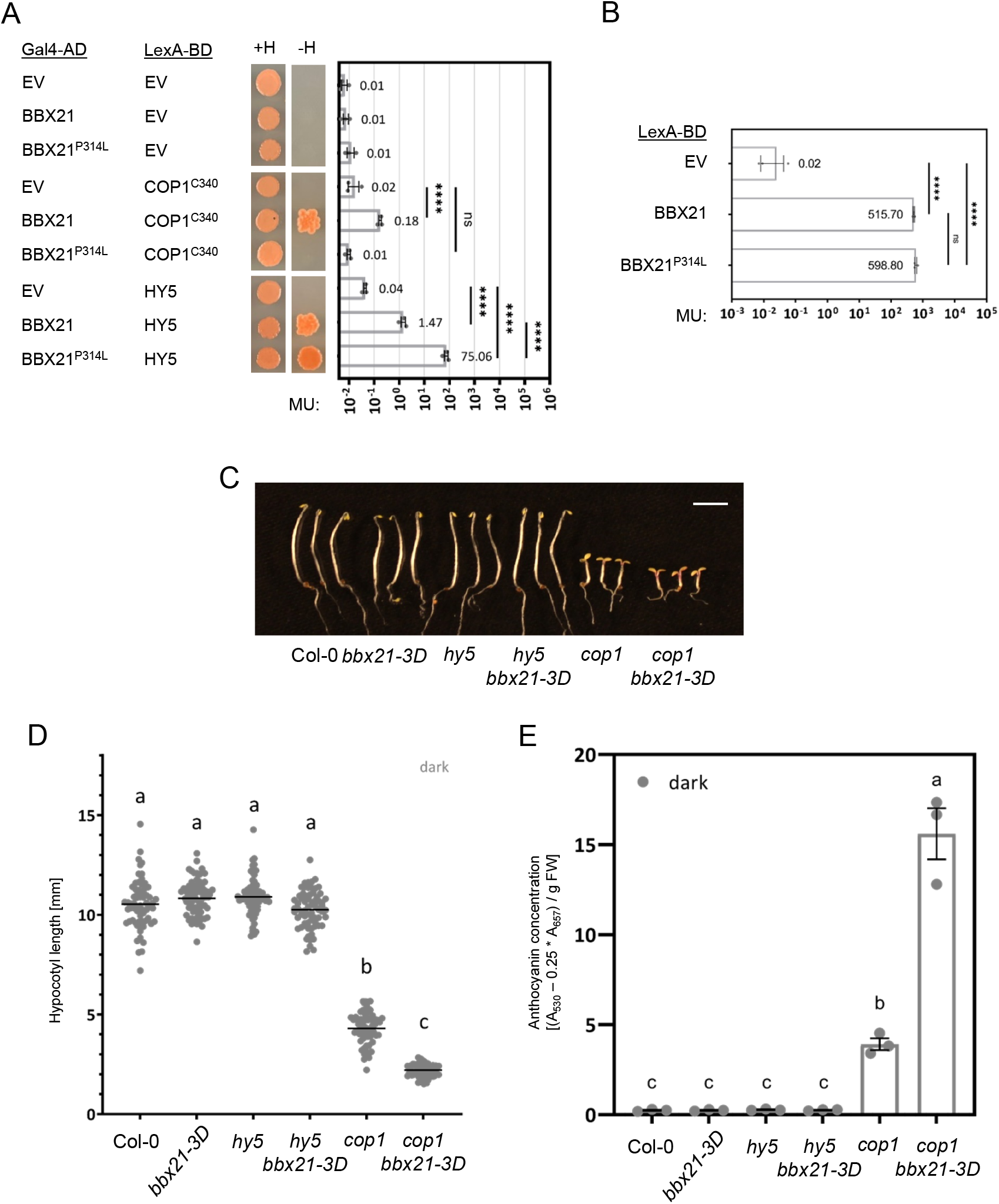
The BBX21^P314L^ variant shows higher activity in a both COP1-dependent and -independent manner. A, Yeast two-hybrid analysis of the interactions of BBX21 and BBX21^P314L^ with COP1^C340^ and HY5. Left: growth assay. Right: quantitative β-galactosidase assay. AD, activation domain; BD, DNA binding domain; +H, +His medium (SD/-Trp/-Leu) as transformation control; -H, selective -His medium (SD/-Trp/-Leu/-His); MU, Miller units; ns, non-significant (P > 0.05); ^****^, P < 0.0001. B, Analysis of the transactivation activity of BBX21 and BBX21^P314L^ fused to the LexA DNA binding domain using a quantitative β-galactosidase assay in yeast. BD, DNA binding domain; MU, Miller units; ns, non-significant (P > 0.05); ^****^, P < 0.0001. C–E, Representative images (C; scale bar indicates 5 mm), quantification of hypocotyl lengths (D; values of independent measurements and means as horizontal lines are shown; *n* > 60), and quantification of anthocyanin concentrations (E; values of independent measurements, means, and SEM are shown; *n* = 3) in wild-type (Col-0), *bbx21-3D, hy5, hy5 bbx21-3D, cop1*, and *cop1 bbx21-3D* seedlings grown in darkness. (D,E) Shared letters indicate no statistically significant difference between the means (P > 0.05).

We further compared the phenotypes of *hy5-215 bbx21-3D* (*hy5 bbx21-3D*) and *cop1-4 bbx21-3D* (*cop1 bbx21-3D*) double mutants to their corresponding single mutants. *cop1 bbx21-3D* showed an enhanced constitutively photomorphogenic phenotype compared to *cop1* (Fig. 2C–E; see also Fig. 3 and Supplemental Fig. S4). This supports the notion that BBX21^P314L^ may exhibit increased activity independent of its protein stabilization through removed COP1-mediated degradation. On the other hand, *hy5 bbx21-3D* showed no aberrant phenotype in darkness, as was observed for *hy5* and *bbx21-3D* (Fig. 2C–E). However, under all light conditions tested, including UV-B (Fig. 3) and monochromatic red and blue light (Supplemental Fig. S4), *hy5* was epistatic to *bbx21-3D*, suggesting that the enhanced photomorphogenic phenotype of *bbx21-3D* is dependent on functional HY5.

**Figure 3.**
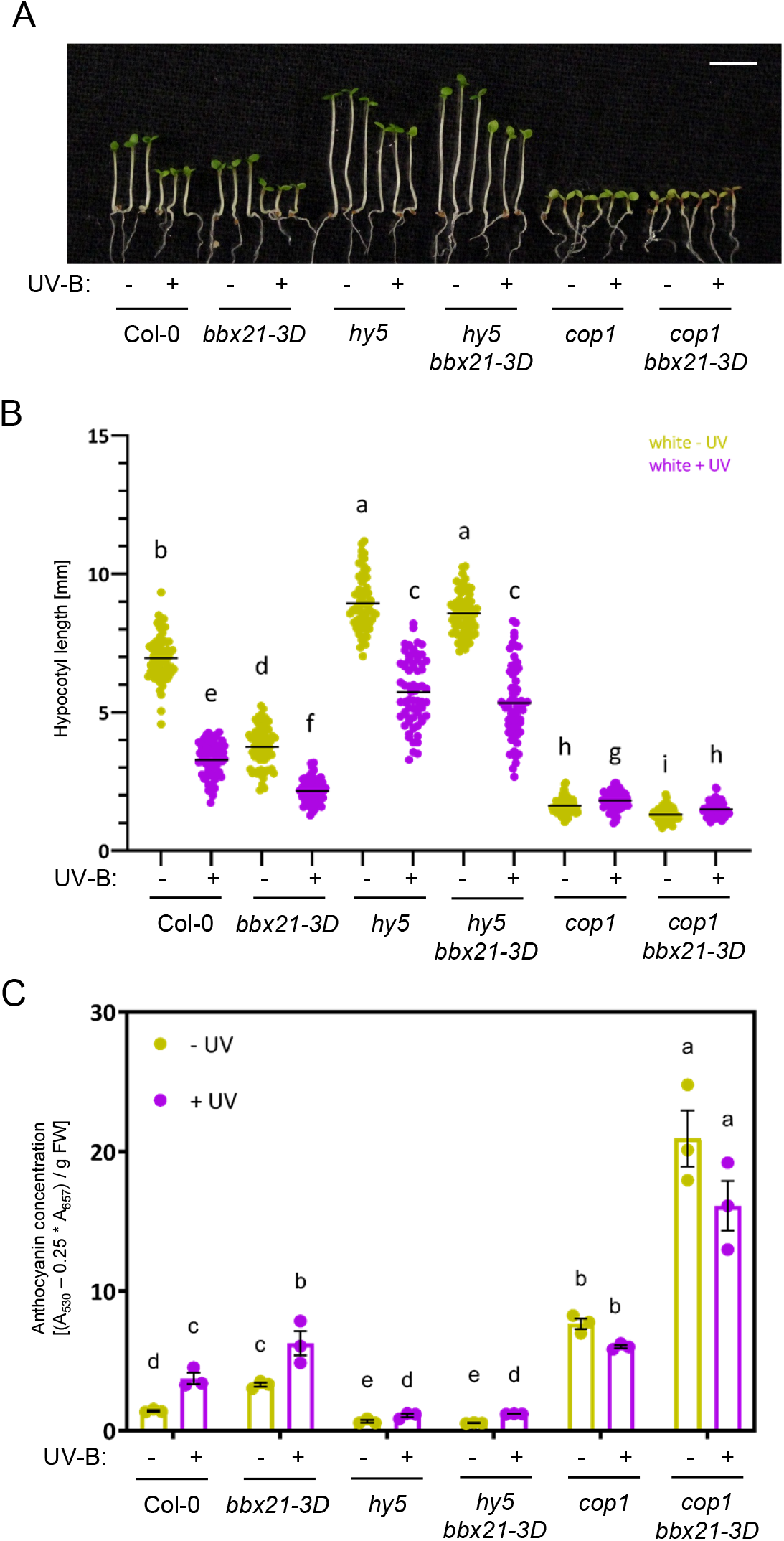
The *bbx21-3D* phenotype requires functional HY5. A–C, Representative images (A; scale bar indicates 5 mm), quantification of hypocotyl lengths (B; values of independent measurements and means as horizontal lines are shown; *n* > 60), and quantification of anthocyanin concentrations (C; values of independent measurements, means, and SEM are shown; *n* = 3) in wild-type (Col-0), *bbx21-3D, hy5, hy5 bbx21-3D, cop1*, and *cop1 bbx21-3D* seedlings grown in weak white light supplemented with UV-B (+), or not (–). (B,C) Shared letters indicate no statistically significant difference between the means (P > 0.05).

### BBX21 and BBX22 are stabilized in response to UV-B

The analysis of *bbx21* null and hypermorphic mutants suggested a role for class IV BBX proteins, which include BBX20 and BBX22, in UV-B signaling. Whereas *BBX20* was apparently not transcriptionally regulated by UV-B, *BBX21* transcript levels were slightly repressed by UV-B (Fig. 4A; see also Supplemental Fig. S2B). By contrast, within 1 h of UV-B exposure, *BBX22* expression induction was observed that was dependent on both UVR8 and HY5 (Fig. 4A,B). Beyond transcriptional regulation, BBX20, BBX21 and BBX22 proteins are post-translationally regulated by COP1 (Chang et al., 2011; Fan et al., 2012; Xu et al., 2016; Job et al., 2018). We thus tested BBX protein levels in lines expressing GFP-tagged BBX20, BBX21, and BBX22 under the constitutive cauliflower mosaic virus (CaMV) 35S promoter. GFP-BBX20 showed constitutive protein levels under UV-B (Fig. 4C), in agreement with the absence of a conserved VP motif in this protein (Supplemental Fig. S5). By contrast, transient stabilization of GFP-BBX21 was detectable after 1 h of UV-B (Fig. 4D). GFP-BBX22 was strongly stabilized in response to UV-B, with immunoblots revealing two specific bands (Fig. 4E), previously reported as representing full-length and truncated forms of GFP-BBX22 (Chang et al., 2011). Full-length GFP-BBX22 levels peaked following approximately 3 h UV-B exposure, whereas the truncated GFP-BBX22 accumulated at least up to 9 h UV-B treatment (Fig. 4E). Using antibodies specifically raised against a BBX22^199–213^ peptide, we observed a similar pattern for endogenous BBX22 in wild type (Fig. 4F). We conclude that BBX21 and BBX22 are transiently stabilized under UV-B, consistent with the notion that their COP1-mediated degradation is relieved once UVR8 binds and represses COP1 activity.

**Figure 4.**
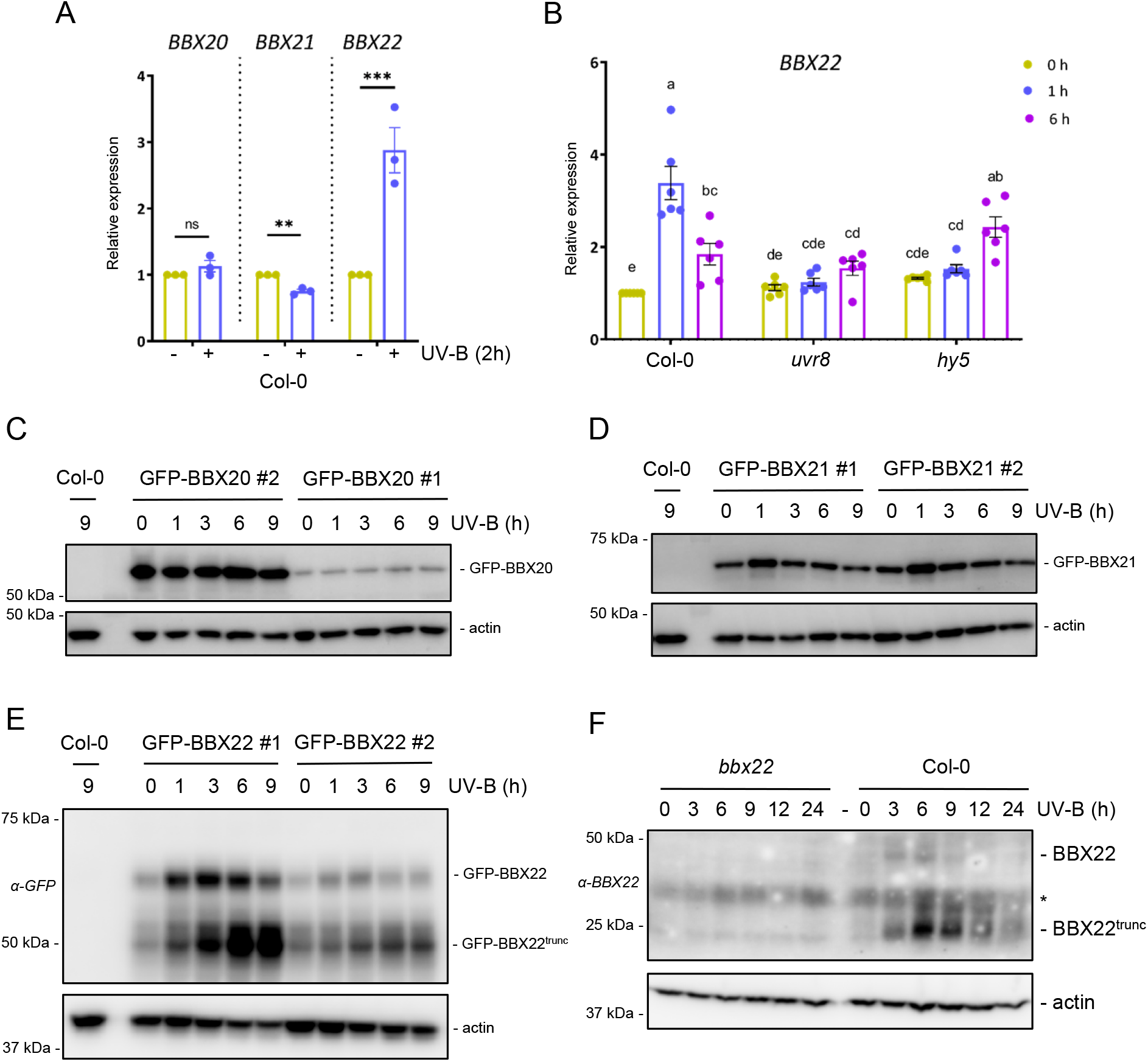
BBX21 and BBX22 are stabilized in response to UV-B. A, RT-qPCR analysis of *BBX20, BBX21*, and *BBX22* expression in 4-d-old wild-type (Col-0) seedlings grown in white light and exposed to 2 h of supplemental UV-B (blue bars), or not (dark yellow bars). B, RT-qPCR analysis of *BBX22* expression in 4-d-old Col-0, *uvr8-12* (*uvr8*), and *hy5* seedlings grown in white light and exposed to 1 h and 6 h of supplemental UV-B, or not (0 h). C–E, Immunoblot analysis of GFP-BBX20 (C), GFP-BBX21 (D), GFP-BBX22 (E), and actin (loading control) levels in two independent lines of each Col-0/Pro_35S_:GFP-BBX20 (GFP-BBX20 #1 and #2), Col-0/Pro_35S_:GFP-BBX21 (GFP-BBX21 #1 and #2), and Col-0/Pro_35S_:GFP-BBX22 (GFP-BBX22 #1 and #2), with Col-0 as negative control. 4-d-old seedlings were treated for 0–9 h with supplemental UV-B, as indicated. F, Immunoblot analysis of endogenous BBX22 and actin (loading control) levels in *bbx22* and Col-0 seedlings treated for 0–24 h with supplemental UV-B, as indicated. ^*^ indicates nonspecific cross-reacting bands. (A,B) Values of independent measurements, means, and SEM are shown (A, *n* = 3; B, *n* = 6). (A) ns, non-significant (P > 0.05); ^**^, P < 0.01; ^***^, P < 0.001. (B) Shared letters indicate no statistically significant difference between the means (P > 0.05).

### HY5 negatively regulates BBX21 and BBX22 accumulation under white light and UV-B

To confirm that the observed BBX protein stabilization was indeed dependent on the UVR8 signaling pathway, we analyzed BBX21 and BBX22 levels in different genetic backgrounds. As expected, GFP-BBX21 and GFP-BBX22 protein stabilization under UV-B was absent in *uvr8-12* null mutants (Fig. 5A,B). This was confirmed for endogenous BBX22 (Fig. 5C). Conversely, in comparison to wild type, endogenous full-length and truncated BBX22 showed strongly enhanced levels in response to UV-B in *uvr8-17D* and *rup1 rup2* (Fig. 5C), which are UV-B hypersensitive mutants due to impaired UVR8 inactivation through redimerization (Gruber et al., 2010; Heijde and Ulm, 2013; Podolec et al., 2021b). Interestingly, in the absence of HY5, strongly enhanced accumulation of GFP-BBX21 was observed before, after 1 h and particularly after 6 h of UV-B treatment (Fig. 5A). This contrasts with the very transient accumulation of GFP-BBX21 in wild type at 1 h of UV-B (Fig. 4D and Fig. 5A). These data suggest that HY5 negatively regulates BBX21 accumulation and is required for the transient nature of BBX21 stabilization under UV-B in wild type. A similar observation was made regarding BBX22 accumulation, where the absence of HY5 allowed the over-accumulation of BBX22 and shifted the peak of full-length BBX22 stabilization to later timepoints (Fig. 5B,D). BBX22 physically interacts with the bZIP domain of HY5 (Datta et al., 2008) and, together with BBX20 and BBX21, forms a transcriptional module that promotes photomorphogenesis (Bursch et al., 2020). We speculated that the repressive effect of HY5 on BBX22 levels may be explained by co-degradation of HY5 and BBX22 by COP1. However, this was not the case, as BBX22 was degraded normally in a *hy5*/Pro_35S_:HY5^ΔN77^ line where the COP1-interacting N-terminus of HY5 is deleted and HY5 is thus constitutively stable (Fig. 5E) (Ang et al., 1998; Osterlund et al., 2000). On the other hand, BBX22 stabilization was strongly repressed in *bbx21-3D* and Pro_35S_:GFP-BBX21 lines (Fig. 5F). Overall, our data suggest that HY5 together with its BBX21 coactivator induce a repressor of BBX22 (and possibly BBX21) protein stability as part of a negative feedback loop included in the photomorphogenic program.

**Figure 5.**
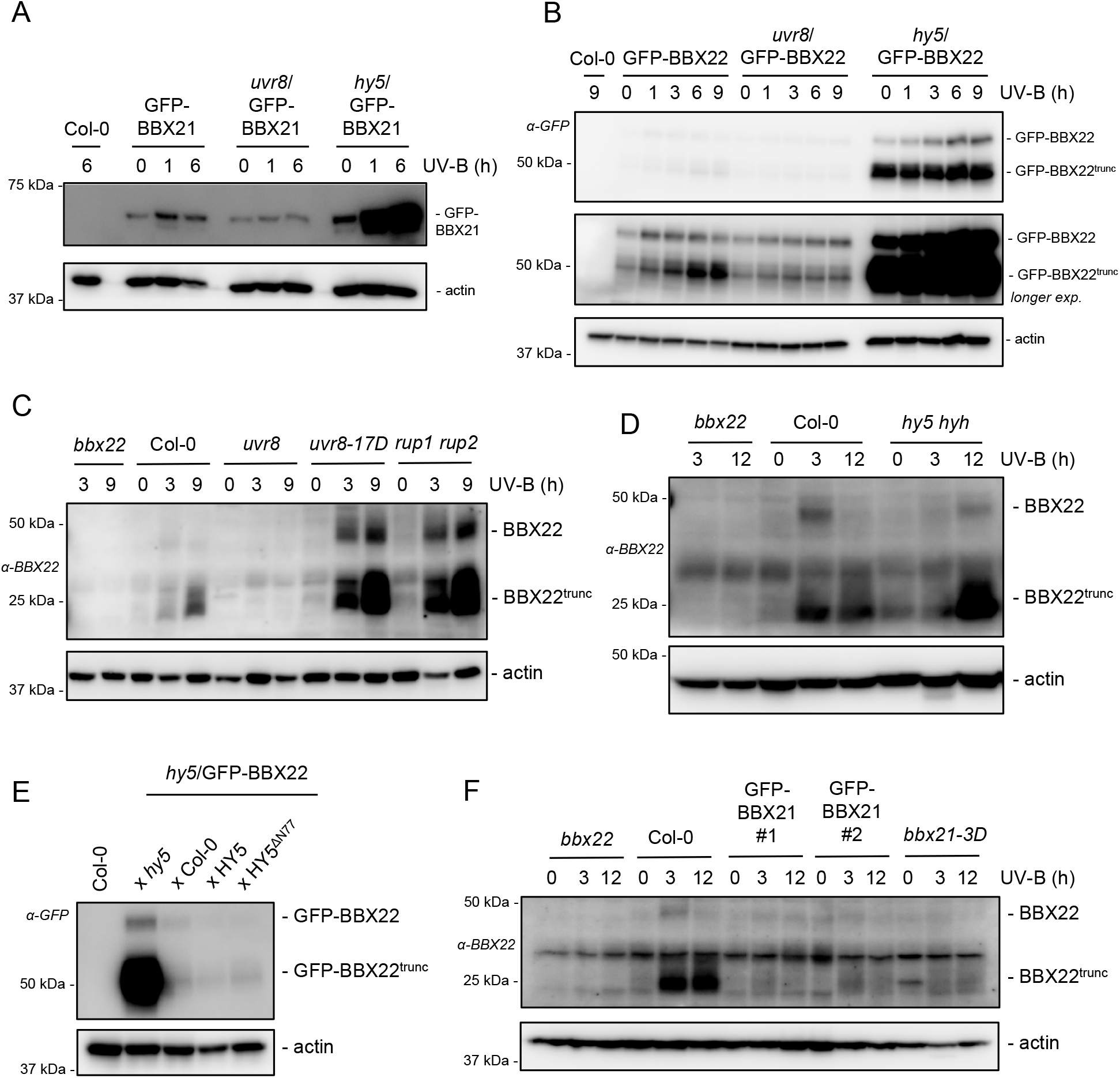
BBX21 and BBX22 accumulation under UV-B is UVR8-dependent and negatively regulated by HY5 activity. A, Immunoblot analysis of GFP-BBX21 and actin (loading control) levels in wild-type (Col-0), Col-0/Pro_35S_:GFP-BBX21 line #2 (GFP-BBX21), *uvr8-12*/Pro_35S_:GFP-BBX21 line #2 (*uvr8*/GFP-BBX21), and *hy5-215*/Pro_35S_:GFP-BBX21 line #2 (*hy5*/GFP-BBX21) seedlings grown for 4 d and treated for 0–6 h with supplemental UV-B, as indicated. B, Immunoblot analysis of GFP-BBX22 and actin (loading control) levels in Col-0, Col-0/Pro_35S_:GFP-BBX22 line #1 (GFP-BBX22), *uvr8-12*/Pro_35S_:GFP-BBX22 line #1 (*uvr8*/GFP-BBX22), and *hy5-215*/Pro_35S_:GFP-BBX22 line #1 (*hy5*/GFP-BBX22) seedlings grown for 4 d and treated for 0– 9 h with supplemental UV-B, as indicated. C, Immunoblot analysis of endogenous BBX22 and actin (loading control) levels in *bbx22*, Col-0, *uvr8-12* (*uvr8*), *uvr8-17D*, and *rup1 rup2* seedlings treated for 0–9 h with supplemental UV-B, as indicated. D, Immunoblot analysis of endogenous BBX22 and actin (loading control) levels in *bbx22*, Col-0, and *hy5 hyh* seedlings treated for 0–12 h with supplemental UV-B, as indicated. E, Immunoblot analysis of GFP-BBX22 and actin (loading control) levels in Col-0, and F1 progeny from crosses of *hy5*/GFP-BBX22 with *hy5*, Col-0, *hy5-215*/Pro_35_S:HY5 line #15 (HY5), and *hy5-215*/Pro_35S_:HY5^ΔN77^ line #27 (HY5^ΔN77^), grown for 4 d in white light. F, Immunoblot analysis of endogenous BBX22 and actin (loading control) levels in *bbx22*, Col-0, Col-0/Pro_35S_:GFP-BBX21 #1 (GFP-BBX21 #1), Col-0/Pro_35S_:GFP-BBX21 #2 (GFP-BBX21 #2), and *bbx21-3D* seedlings treated for 0–12 h with supplemental UV-B, as indicated.

### BBX20, BBX21, and BBX22 promote UVR8- and HY5-dependent responses

To determine whether BBX proteins contribute to UVR8- and HY5-dependent, UV-B-induced photomorphogenic responses, we tested the phenotypes of combinatorial mutants, including a *bbx20 bbx21 bbx22* triple mutant. As shown previously (Favory et al., 2009; Tavridou et al., 2020), *uvr8* mutant seedlings were strongly impaired in UV-B-induced hypocotyl growth inhibition, whereas *hy5* showed longer hypocotyls under both –UV-B and +UV-B conditions, with reduced inhibition of hypocotyl elongation under UV-B (Fig. 6A,B and Supplemental Fig. S6A). The *bbx20 bbx21 bbx22* triple mutant displayed longer hypocotyls in –UV-B conditions, similar to *hy5*, and an intermediate hypocotyl length between that of wild type and *hy5* under UV-B (Fig. 6A,B and Supplemental Fig. S6A). These data indicate that BBX proteins partially regulate the HY5-dependent inhibition of hypocotyl elongation under UV-B. On the other hand, anthocyanin accumulation was strongly compromised in the *bbx20 bbx21 bbx22* triple mutant, resembling *hy5* (Fig. 6C). Flavonol profiling by high performance thin layer chromatography (HPTLC) further revealed that *bbx20 bbx21 bbx22* was strongly impaired in the UV-B-dependent accumulation of flavonol glycosides (Fig. 6D). It is of note that the *hy5 bbx20 bbx21 bbx22* quadruple mutant showed a weak additive phenotype for both hypocotyl length and pigment accumulation, suggesting that a HY5-independent activity of BBX proteins exists in addition to the above described BBX-independent activity of HY5 (Fig. 6A–C). Related to pigment biosynthesis, UV-B-induced CHS protein accumulation was impaired but not completely abolished in *bbx20 bbx21 bbx22*, whereas it was undetectable in *hy5* (Fig. 6E and Supplemental Fig. S6B), suggesting there are additional important regulators of HY5 activity besides BBX20, BBX21, and BBX22. In agreement with reduced CHS accumulation and impaired pigment biosynthesis, *CHS* transcript did not accumulate to high levels under prolonged UV-B conditions in *bbx20 bbx21 bbx22*, similar to *hy5* and unlike in UV-B-exposed wild-type seedlings (Fig. 6F).

**Figure 6.**
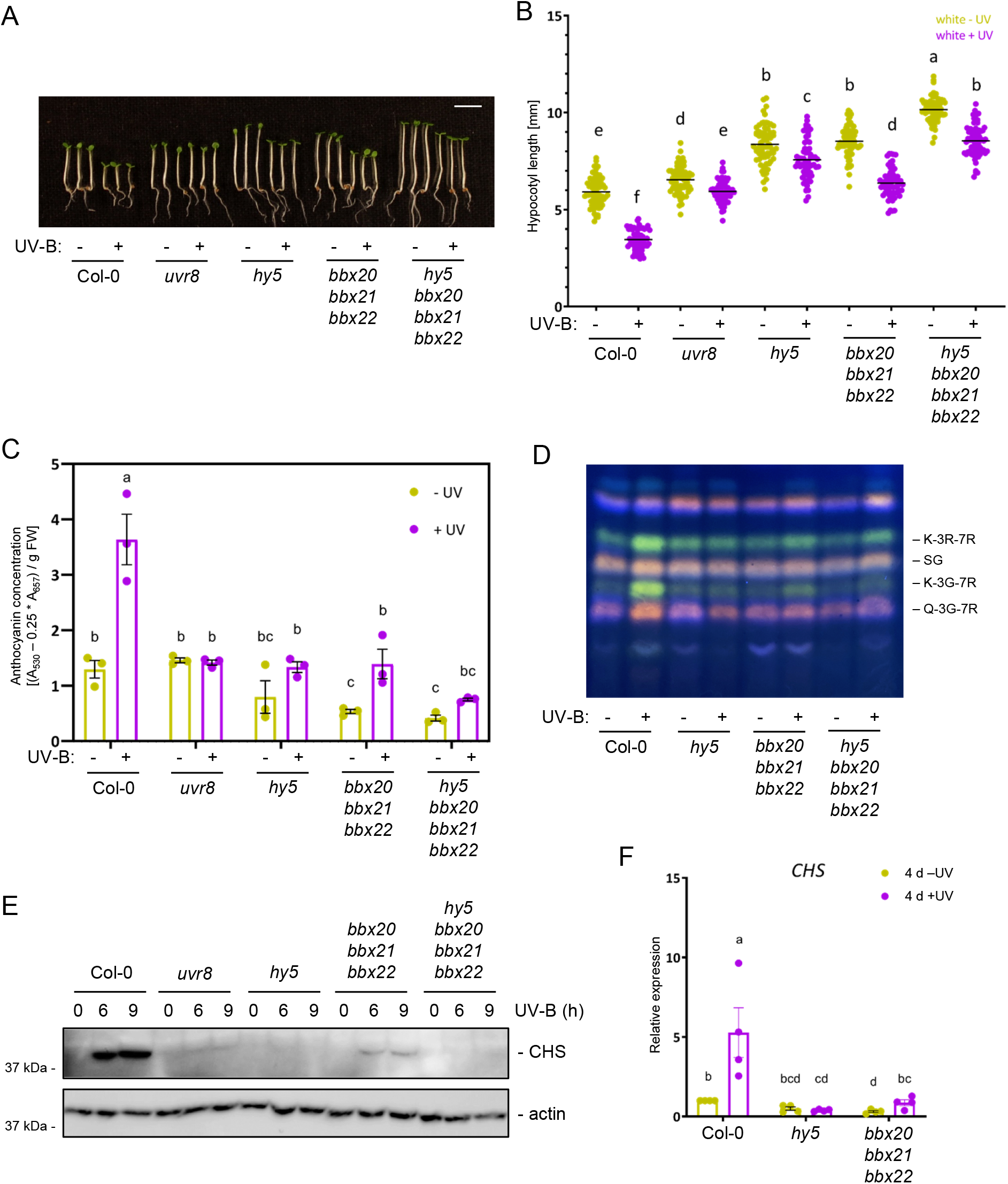
BBX proteins are necessary for hypocotyl growth inhibition and pigment accumulation under UV-B. A, Representative images of wild-type (Col-0), *uvr8-12* (*uvr8*), *hy5-215* (*hy5*), *bbx20 bbx21 bbx22*, and *hy5 bbx20 bbx21 bbx22* mutant seedlings in white light supplemented (+) or not (–) with UV-B. Scale bar indicates 5 mm. B, Quantification of hypocotyl length of seedlings as shown in (A). C, Anthocyanin concentrations in seedlings as shown in (A). D, HPTLC analysis of the flavonol glycoside levels in seedlings as described in (A) grown in white light or white light supplemented with UV-B. K-3R-7R, kaempferol-3-O-rhamnoside-7-O-rhamnoside; SG, sinapoyl glucose; K-3G-7R, kaempferol-3-O-glucoside-7-O-rhamnoside; Q-3G-7R, quercetin-3-O-glucoside-7-O-rhamnoside. E, Immunoblot analysis of CHS and actin (loading control) levels in seedlings as described in (A) that were exposed for 0–9 h to supplemental UV-B, as indicated. F, RT-qPCR analysis of *CHS* expression in Col-0, *hy5*, and *bbx20 bbx21 bbx22* seedlings grown for 4 d in white light or white light supplemented with UV-B. (B) Values of independent measurements and means as horizontal lines are shown (*n* > 60). (C,F) Values of independent measurements, means, and SEM are shown (C, *n* = 3; F, *n* = 4). (B,C,F) Shared letters indicate no statistically significant difference between the means (P > 0.05).

### BBX20, BBX21, and BBX22 promote the expression of some genes under prolonged UV-B but not their short-term induction after UV-B exposure

BBX proteins regulate *HY5* transcription (Xu et al., 2016; Bursch et al., 2020); however, both *HY5* transcript and HY5 protein accumulation after UV-B exposure were not strongly affected in the *bbx20 bbx21 bbx22* triple mutant (Fig. 7A,B). HY5-dependent induction of UV-B marker genes such as *RUP2, CHS*, and *ELIP2* was only weakly, if at all, affected in *bbx20 bbx21 bbx22* in response to UV-B (Fig. 7C–E). Overall, the induction of transcripts was largely independent of BBX20, BBX21, and BBX22, and the residual induction in *bbx20 bbx21 bbx22* was HY5 dependent (Fig. 7B–E). This was in stark contrast to the impaired induction of *CHS* after 4 d of continuous UV-B conditions (Fig. 6F). Thus, we checked the expression of other marker genes in continuous –UV-B and +UV-B conditions. Interestingly, in *bbx20 bbx21 bbx22, F3H* induction under prolonged UV-B was impaired as well, whereas *MYB12* and *ELIP2* were only partially affected (Fig. 8A–C). By contrast, whereas *RUP1* and *RUP2* UV-B induction maintained strong dependence on HY5, *RUP1* and *RUP2* transcripts accumulated normally under prolonged UV-B in *bbx20 bbx21 bbx22* (Fig. 8D,E). In summary, our data suggest that BBX20, BBX21, and BBX22 redundantly promote pigment biosynthesis and the accumulation of some transcripts, such as *CHS* and *F3H*, under prolonged UV-B. However, BBX20, BBX21, and BBX22 do not seem to regulate the early induction of *HY5* and other UV-B marker genes immediately after UV-B exposure; and they play a partial role in the HY5-dependent inhibition of hypocotyl elongation.

**Figure 7.**
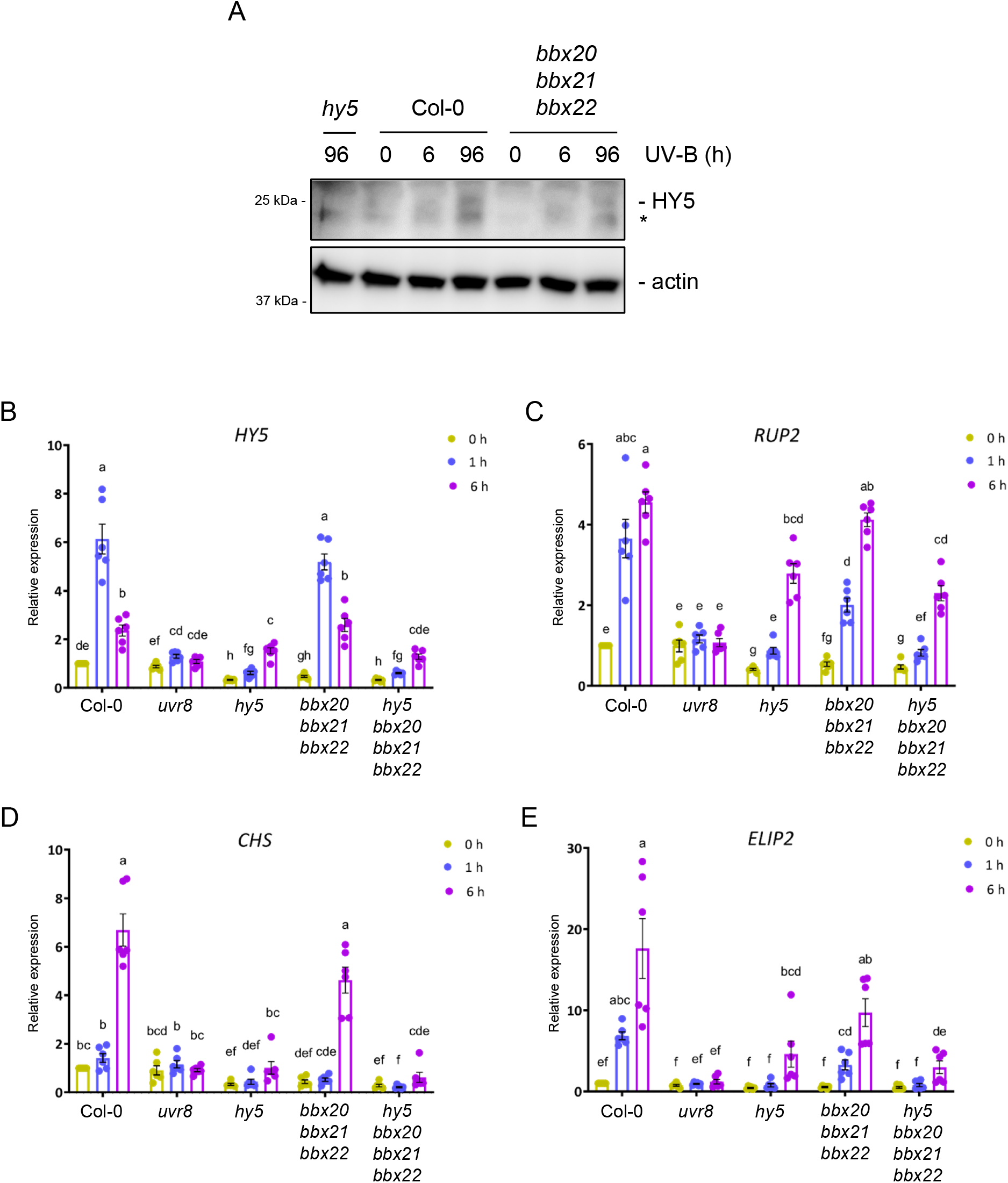
BBX proteins play a minor role in the short-term induction of UV-B marker genes. A, Immunoblot analysis of HY5 and actin (loading control) levels in wild-type (Col-0), *hy5*, and *bbx20 bbx21 bbx22* seedlings exposed to 6 h and 96 h of supplemental UV-B, or not (0). B–E, RT-qPCR analysis of *HY5* (B), *RUP2* (C), *CHS* (D), and *ELIP2* (E) expression in 4-d-old Col-0, *uvr8, hy5, bbx20 bbx21 bbx22*, and *hy5 bbx20 bbx21 bbx22* seedlings grown in white light and exposed to 1 h and 6 h of supplemental UV-B, or not (0 h). Values of independent measurements, means, and SEM are shown (*n* = 6); shared letters indicate no statistically significant difference between the means (P > 0.05).

**Figure 8.**
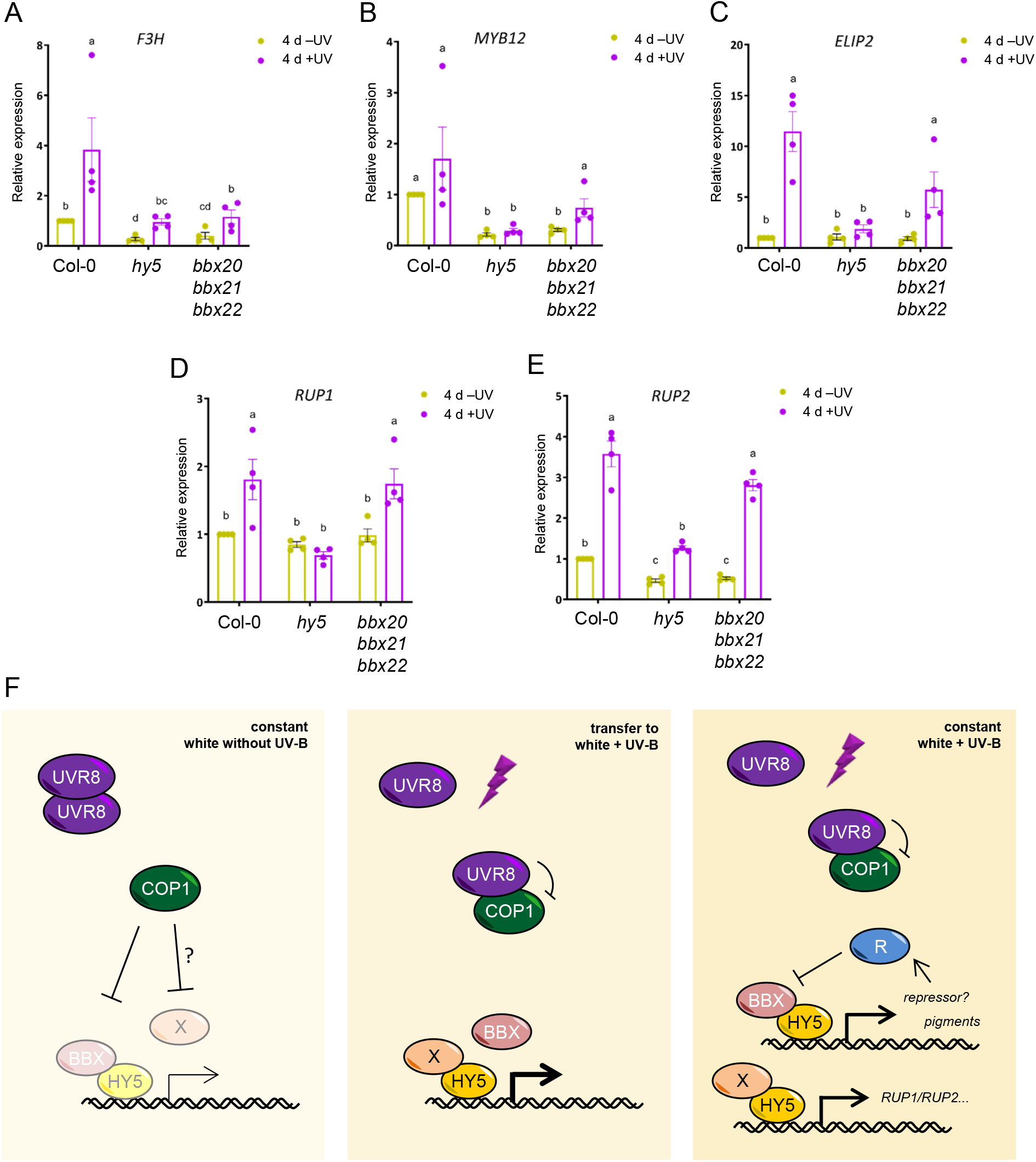
BBX proteins are necessary for the expression of some but not all marker genes in white light conditions constantly supplemented with UV-B. A–E, RT-qPCR analysis *F3H* (A), *MYB12* (B), *ELIP2* (C), *RUP1* (D), and *RUP2* (E) expression in wild-type (Col-0), *hy5*, and *bbx20 bbx21 bbx22* seedlings grown for 4 d in white light or white light supplemented with UV-B. Values of independent measurements, means, and SEM are shown (*n* = 4); shared letters indicate no statistically significant difference between the means (P > 0.05). F, Working model for a role of BBX proteins in UV-B photomorphogenesis. Left – Under weak white light conditions devoid of UV-B, UVR8 is inactive and COP1 targets downstream factors, including HY5, BBX proteins, and possibly a hypothetical factor X, for degradation. A small residual pool of HY5 and BBX proteins are responsible for the low basal expression of light-induced marker genes and a weak inhibition of hypocotyl elongation. Middle – Upon exposure of seedlings to UV-B, UVR8 represses COP1 activity. This results in the accumulation of COP1 targets: HY5, BBX proteins, and factor X. HY5 increasingly binds promoters of light-responsive genes, and factor X provides it with transcriptional activity, so that the expression of marker genes is strongly induced. BBX proteins play a minor role in this transcriptional induction. Right – In prolonged UV-B conditions, COP1 remains repressed by UVR8, but this is dampened by negative feedback on photoreceptor activity through RUP1/RUP2. HY5 remains highly stable, but BBX protein levels are repressed through the action of a hypothetical repressor (R) that is induced as a part of HY5/BBX-mediated photomorphogenesis. BBX proteins play a role as HY5 coactivators in allowing the sustained expression of genes involved in pigment biosynthesis, whereas other UV-B marker genes are induced by HY5 in a BBX-independent manner. Hypocotyl length inhibition is controlled by both BBX proteins and factor X.

## DISCUSSION

Upon UV-B perception, nuclear UVR8 promotes photomorphogenesis and acclimation by directly or indirectly modulating the activity of several transcriptional regulators (Favory et al., 2009; Qian et al., 2016; Yin et al., 2016; Liang et al., 2018; Yang et al., 2018; Liang et al., 2019; Qian et al., 2020; Podolec et al., 2021a). These include the bZIP transcriptional regulators HY5 and HYH that control the expression of many genes downstream of UVR8 and both the cryptochrome and phytochrome pathways (Ang et al., 1998; Holm et al., 2002; Ulm et al., 2004; Brown et al., 2005; Oravecz et al., 2006; Stracke et al., 2010; Burko et al., 2020). These photoreceptors inhibit the activity of the COP1/SPA E3 ubiquitin ligase complex, allowing accumulation of HY5 and HYH followed by induction of the photomorphogenic program in the light (Hoecker, 2017; Podolec and Ulm, 2018). Despite its important and broad function as transcriptional regulator, HY5 itself lacks a transcriptional activation domain (Ang et al., 1998; Stracke et al., 2010; Burko et al., 2020). Recently, BBX20, BBX21 and BBX22 were characterized as HY5 coregulators that provide transactivational activity, allowing transcriptional gene activation in response to both red light and the butenolide molecule karrikin (Bursch et al., 2020; Bursch et al., 2021).

Here, we uncovered a novel, hypermorphic *bbx21* allele in a genetic screen for mutants with enhanced UV-B responses. BBX21 is a promoter of photomorphogenesis that is targeted by COP1 for degradation (Xu et al., 2016). We identified a *bbx21-3D* mutant containing a P314L mutation, which functionally abolished the COP1 interaction domain of BBX21. This novel VP-mutated allele of a COP1 substrate allowed us to test the functional relevance of a VP motif in an endogenous context. The *bbx21-3D* phenotype was largely consistent with that of previously published *BBX21* overexpression lines (Xu et al., 2016; Job et al., 2018), supporting the assumption that BBX21^P314L^ is post-translationally stabilized and thus present at elevated levels in the mutant background, as expected for a BBX21 VP mutant (Podolec and Ulm, 2018; Lau et al., 2019; Bursch et al., 2020). Unfortunately, we were not able to raise specific anti-BBX21 antibodies to probe directly for endogenous BBX21^P314L^ levels compared to wild-type BBX21. Interestingly, however, the *bbx21-3D* mutation enhanced the *cop1-4* mutant phenotype, although no interaction between BBX21 and COP1 is expected to occur in this genetic context since the corresponding COP1^N282^ protein expressed in *cop1-4* lacks the VP-interacting WD40 repeat domain (McNellis et al., 1994; Lau et al., 2019). These data suggest that BBX21^P314L^ also exhibits elevated activity independently of its impaired interaction with and regulation by COP1. The additive phenotype of *cop1* and *bbx21-3D* could be due to a stronger HY5–BBX21^P314L^ interaction, as our yeast interaction data suggested, and thus increased activity of BBX21 as a HY5 coactivator (Bursch et al., 2020).

We found similarities but also interesting differences in the role of BBX proteins in UV-B signaling compared to that under visible light. Regarding hypocotyl elongation phenotype, *hy5* and *bbx20 bbx21 bbx22* were comparably elongated in constant white light devoid of UV-B, but differed under constant white light supplemented with UV-B, wherein BBX proteins accounted for only a part of the HY5-dependent response. Interestingly, and similar to our observations, the HY5-dependent hypocotyl response of seedlings during karrikin signaling relies only partially on BBX20 and BBX21 (Bursch et al., 2021). Furthermore, the BBX-dependent transcriptome accounted only for a fraction of HY5-regulated transcription under red light, suggesting that additional HY5 coregulators are indeed involved in some responses (Bursch et al., 2020). In contrast to the hypocotyl elongation phenotype, BBX proteins were crucial for UV-B-dependent biosynthesis of phenylpropanoid pigments (flavonols, anthocyanins). Intriguingly, when analyzing the molecular basis of this phenotype, we found that gene expression involved in phenylpropanoid biosynthesis (*CHS, F3H*) was impaired under prolonged and constant UV-B, but the induction of early UV-B marker genes within 6 h of UV-B exposure was almost completely intact. Similarly, CHS protein accumulation after UV-B exposure was decreased but not abolished in the *bbx20 bbx21 bbx22* triple mutant. These data suggest that early and transient HY5 responses to UV-B are BBX20, BBX21, and BBX22 independent, whereas for long-term responses such as pigment biosynthesis, HY5 relies on these BBX proteins as transcriptional coactivators. It is of note that the long-term accumulation of some transcripts, such as *RUP1* and *RUP2*, was also BBX-independent.

BBX21 and BBX22 were post-transcriptionally stabilized under UV-B in a UVR8-dependent manner. This is consistent with their status as COP1 substrates (Chang et al., 2011; Xu et al., 2016; this work) and how active UVR8 directly inhibits COP1 activity by competitively blocking the COP1 substrate binding site (Lau et al., 2019; Podolec et al., 2021a). Consistent with previous literature on visible light responses (Chang et al., 2011; Xu et al., 2016; Job et al., 2018), we observed a transient stabilization of BBX21 and BBX22, with protein levels decreasing after a few hours of UV-B exposure. This transient stabilization could be linked to attenuation of UVR8 signaling through the RUP1- and RUP2-mediated negative feedback loop (Gruber et al., 2010; Heijde and Ulm, 2013; Ren et al., 2019; Podolec et al., 2021b), as supported by extended BBX22 accumulation in *uvr8-17D* and *rup1 rup2* backgrounds. Moreover, our data indicate the existence of an additional negative regulator of BBX21 and BBX22 stability. This repressor seems to be linked to HY5 activity, as suggested by the overaccumulation of BBX21 and BBX22 in *hy5* backgrounds and as already reported for BBX22 under visible light (Chang et al., 2011). Conversely, plant lines with enhanced photomorphogenesis (such as *bbx21-3D* and BBX21 overexpression lines) show lower levels of BBX22 accumulation. Collectively, our data point to an uncharacterized negative feedback mechanism that attenuates photomorphogenesis induced by the HY5/BBX transcriptional module.

We conclude that BBX20, BBX21, and BBX22 play important roles as HY5 coactivators in inducing UV-B responses such as pigment biosynthesis, and to a lesser degree inhibition of hypocotyl elongation, mainly in constant UV-B conditions (for our working model, see Fig. 8F). Short-term responses on the other hand (few hours after UV-B exposure) seem to be mostly independent of these BBX proteins, as seen by the normal transcriptional induction of HY5-dependent UV-B marker genes in *bbx20 bbx21 bbx22*. This further indicates the existence of a specific HY5 coactivator, or multiple thereof, under UV-B, the activation of which is likely crucial for early gene activation upon UV-B reception and signaling by the UVR8 photoreceptor. Whether COP1 regulates further HY5 coregulators and whether they are members of the large BBX family remains to be determined.

## MATERIALS AND METHODS

### Plant materials

All lines used in this study are in the Arabidopsis (*Arabidopsis thaliana*) Columbia (Col-0) background. The following lines have been described previously: *rup1 rup2* (Gruber et al., 2010), *cop1-4* (Deng et al., 1992), *uvr8-12, uvr8-17D* (Podolec et al., 2021b), *hy5-215* (Oyama et al., 1997), *hy5-215 hyh* (Zoulias et al., 2020), *bbx21-1* (Datta et al., 2007), *bbx22-1* (Chang et al., 2008), *bbx20-1, bbx21-1 bbx22-1, bbx20-1 bbx21-1 bbx22-1, hy5-215 bbx20-1 bbx21-1 bbx22-1*, Col-0/Pro_35S_:GFP-BBX20 #1 and #2, *hy5-215*/Pro_35S_:HY5^ΔN77^ #27, *hy5-215*/Pro_35S_:GFP-BBX21 #2 (Bursch et al., 2020), *bbx20-1 bbx21-1* (Bursch et al., 2021), Col-0/Pro_35S_:GFP-BBX21 #1 and #2, *hy5-215*/Pro_35S_:HY5 #15 (Job et al., 2018). Pro_35S_:GFP-BBX22 lines #1 and #2 were generated in the Col-0 background using a pB7WGF2 vector (Karimi et al., 2002) in which the *BBX22* CDS was inserted after cloning into pDONR221 (primers BBX22_attB1_Fw and BBX22_attB2_Rv, Table S1) using Gateway technology (ThermoFisher). *bbx21-3D* was identified in a forward genetic screen based on hypocotyl length under UV-B of an EMS mutagenized Col-0 population (Podolec et al., 2021b), and was backcrossed three times. Combinatorial mutant lines *bbx20-1 bbx22-1, hy5-215 bbx21-3D, cop1-4 bbx21-3D, uvr8-12*/Pro_35S_:GFP-BBX21 #2, *uvr8-12*/Pro_35S_:GFP-BBX22 #1, and *hy5-215*/Pro_35S_:GFP-BBX22 #1 were generated by crossing and genotyped by PCR and sequencing.

### Generation of *BBX21* CRISPR/Cas9-mutated lines

The CRISPR/Cas9 system was used to delete the *BBX21* C-terminus in wild type and to knockout *BBX21* in wild type and *bbx21-3D*. The sgRNA directed against the *BBX21* sequence were inserted into the pHEE401E vector (Wang et al., 2015) using overlapping complementary oligos (Supplemental Table S1). Arabidopsis plants were then transformed using the floral-dip method (Clough and Bent, 1998). Several independent transgenic events were selected and the *BBX21* locus was sequenced in T2 to identify lines with desired mutations.

### Growth conditions and light treatments

Seeds were surface sterilized using chlorine gas or ethanol and sown on ½ MS (Duchefa) agar medium supplemented with 1% (w/v) sucrose (except for experiments in monochromatic red/blue light, which were done on media without sucrose). Plates were left for 2 d in the dark at 4°C for stratification. For growth in darkness, plates were exposed for 6 h to ∼60 µmol m^-2^ s^-1^ of white light and then transferred to darkness at 22°C for 4 d. For light treatments, plates were grown for 4 d at 22°C in the following light conditions: weak white light (3.6 µmol m^-2^ s^-1^, Osram L18W/30 tubes) supplemented or not with narrowband UV-B (1.5 µmol m^-2^ s^-1^, Philips TL20W/01RS tubes), monochromatic red light (150 µmol m^-2^ s^-1^, floraLEDs in CLF Plant Climatics cabinet), monochromatic blue light (50 µmol m^-2^ s^-1^, floraLEDs in CLF Plant Climatics cabinet). For UV-B stress treatment, plates were irradiated with broadband UV-B (21 µmol m^-2^ s^-1^, Philips TL20W/12RS tubes).

### Hypocotyl length measurements

Hypocotyl length was determined as described previously (Podolec et al., 2021b). In short, seedlings were grown for 4 d in the appropriate condition and approximately 60 seedlings for each genotype/condition were aligned on an agar plate and scanned. Individual hypocotyls were measured using the NeuronJ plugin of ImageJ (Meijering et al., 2004).

### Extraction and quantification of anthocyanins

Anthocyanins were quantified as described previously (Yin et al., 2012). Seedlings were grown for 4 d in the appropriate condition and approximately 50 mg of seedlings were collected for each genotype and condition. Samples were frozen in liquid nitrogen, ground, and 250 µl of extraction buffer (99% [v/v] methanol, 1% [v/v] HCl) was added to the samples. Extraction was performed for at least 1 h by incubating the samples on a rotary shaker at 4°C. After centrifugation for 5 min, the absorbance of 150 µl of the clear supernatant was measured at 530 nm and 655 nm. The following formula was used to calculate the relative quantity of anthocyanins: (A_530_ - 0.25 ^*^ A_655_) / seedling mass (in mg).

### Extraction and visualization of flavonoids

High-performance thin layer chromatography (HPTLC) was used to analyze the flavonol profile as described previously (Podolec et al., 2021b). In short, exactly 50 mg of seedlings were collected for each genotype and condition, samples were frozen and then ground. 100 µl of extraction buffer (80% [v/v] methanol) was added per sample, which were then incubated for 10 min at 70°C on a shaker. After a 5 min centrifugation at 14,000 rpm, supernatants were collected and 10 µl was loaded on silica HPTLC plates. The extracts were then separated for approximately 45 min using a mobile phase (5 ml ethyl acetate, 600 µl formic acid, 600 µl acetic acid glacial, 1.3 ml water). After migration, the plate was dried and sprayed with 1% (w/v) diphenylboric acid 2-aminoethylester (DPBA, Roth) solution in 80% (v/v) methanol. The plate was exposed under a 365-nm UV-A lamp to reveal the flavonoid profile.

### Measurements of photosynthetic efficiency

Maximum quantum efficiency of photosystem II was measured after dark-adapting plants for 5 min using a Fluorcam (Photon Systems Instruments) with blue (470 nm) LEDs, and was calculated as Fv/Fm = (Fm – Fo) / Fm, where Fm is the maximal fluorescence and Fo the minimal fluorescence in the dark-adapted state.

### Protein extraction and immunoblot assays

For all immunoblot assays that include (GFP-)BBX levels, a previously described buffer (Job et al., 2018) was used: 50 mM Tris-HCl pH 7.5, 150 mM NaCl, 1% (w/v) sodium deoxycholate, 0.5% (v/v) Triton X100, 1 mM DTT, 50 µM MG132 (Sigma), 50 µM ALLN (VWR), and 50 µM Protease Inhibitor Cocktail (Sigma). For HY5 immunoblots, a previously described buffer (Oravecz et al., 2006) was used: 0.1 M Tris-HCl pH 8.0, 50 µM EDTA, 0.25 M NaCl, 0.7% (w/v) SDS, 10 mM NaF, 15 mM β-glycerolphosphate, 15 mM p-nitrophenyl phosphate, a Complete EDTA-free Protease Inhibitor Cocktail tablet (Roche), and 1 mM DTT. For CHS immunoblot assays, a previously described phosphate buffer (Arongaus et al., 2018) was used: 50 mM Na-phosphate pH 7.4, 150 mM NaCl, 10% (v/v) glycerol, 5 mM EDTA, 0.1% (v/v) Triton X-100, 1 mM DTT, 2 mM Na_3_VO4, 2 mM NaF, 1% (v/v) Protease Inhibitor Cocktail (Sigma), and 50 µM of MG132 (Sigma).

In all cases except for HY5 immunoblots, samples were harvested, frozen, ground, and mixed with extraction buffer before centrifugation for 25 min at 12,000 rpm, 4°C. The clear supernatants were collected and protein concentration was determined using the Bradford-based Bio-Rad protein assay (Bio-Rad). Samples were then denatured and separated using SDS-PAGE. Proteins were transferred on PVDF membranes (Roth) for 7 min at 20 V using the iBlot dry blotting system (Thermo Fisher Scientific), before blocking in TBS-T with milk.

For HY5 immunoblots, proteins were extracted as described above and wet-transferred (transfer buffer: 25 mM Tris; 192 mM glycine; 20% [v/v] ethanol) on PVDF membranes (Roth). Membranes were rinsed once with water, once with TBS-T, then dried for 15 min in a speed-vac at 65°C and kept for 5 d at room temperature in the dark. Subsequent incubation steps were performed in TBS-T without milk.

The following primary antibodies were used: anti-CHS (sc-12620, Santa Cruz Biotechnology), anti-GFP (Living Colors® A.v. Monoclonal Antibody, JL-8; Clontech), anti-HY5 (Oravecz et al., 2006), anti-actin (A0480, Sigma-Aldrich), anti-BBX22^(199–213)^ (Eurogentec, raised in rabbits against the peptide C+DQSYEYMENNGSSKT and affinity purified). Corresponding horseradish peroxidase-conjugated anti-rabbit (for anti-HY5 and anti-BBX22), anti-mouse (for anti-GFP and anti-actin), and anti-goat (for anti-CHS) immunoglobulins (Dako) were used as secondary antibodies. Signal was revealed using the ECL Select Western Blotting Detection Reagent (GE Healthcare) and analyzed using an Amersham Imager 680 camera system (GE Healthcare).

### Yeast two-hybrid assays

Yeast two-hybrid assays were performed after transforming the L40 strain (Vojtek and Hollenberg, 1995) using the lithium-acetate protocol (Gietz, 2014). *BBX21* and *BBX21*^*P314L*^ coding sequences were inserted in frame with the Gal4 activation domain (Gal4-AD) into pGADT7-GW (Marrocco et al., 2006) and *COP1*^*C340*^ and *HY5* were inserted in frame with the LexA DNA binding domain (LexA-DB) into pBTM116-D9-GW (Stelzl et al., 2005) using Gateway cloning. To test for intrinsic transcriptional activation potential, *BBX21* and *BBX21*^*P314L*^ were inserted in frame with LexA-DB into pBTM116-D9-GW (Stelzl et al., 2005) using Gateway cloning. Multiple colonies from each transformation were mixed, spotted, and grown on vector-selection medium (SD/-Trp/- Leu; SD/-Trp for testing transcriptional activation potential). For growth assays, -His selective medium for interactions (SD/-Trp/-Leu/-His, Formedium) was used. For quantitative β-galactosidase assays, yeast cells were grown for 2 d on vector-selection plates, collected and the enzymatic assay was performed using red-β-D-galactopyranoside (CPRG, Roche Applied Science) as substrate as described (Yeast Protocols Handbook, Clontech).

### Gene expression analysis

To determine transcript levels by reverse transcription quantitative PCR (RT-qPCR), RNA was extracted using the ReliaPrep RNA Tissue Miniprep System kit (Promega) and treated with DNase according to the manufacturer’s instructions. cDNA synthesis was performed using the TaqMan reverse transcription kit (Applied Biosystems), with a 1:1 mix of oligo-dT and random hexamer primers. The qPCR reaction was performed using the PowerUp SYBR Green Master Mix reagents (Applied Biosystems) on a QuantStudio™ 5 System (Thermo Fisher Scientific) following the recommended protocol, using primers listed in Supplemental Table S1. The ΔΔCt method (Livak and Schmittgen, 2001) was used to calculate expression values, with *PP2A* as a reference gene. Each experiment was done by combining 3–6 independently made biological replicates. All expression values were normalized within each biological replicate against the untreated wild type that was set to 1.

### Statistical analysis

Statistical analyses were done using the GraphPad Prism software (v. 9.2.0). One- or two-way ANOVAs were performed on log-transformed or non-transformed data, followed by Tukey’s test for multiple comparisons to distinguish statistically different groups (P > 0.05) using different letters. For the analysis of Fv/Fm (Fig. 1G), a repeated measures ANOVA was used. For pairwise comparisons (Fig. 4A), unpaired two-tailed *t*-tests were used and P values were represented.

### Accession numbers

Sequence data from this work can be found in the Arabidopsis Genome Initiative or GenBank/EMBL databases under the following accession numbers: AT4G39070 (BBX20), AT1G75540 (BBX21), AT1G78600 (BBX22), AT5G13930 (CHS), AT3G51240 (F3H), AT2G32950 (COP1), AT5G11260 (HY5), AT3G17609 (HYH), AT2G47460 (MYB12), AT4G14690 (ELIP2), AT5G52250 (RUP1), AT5G23730 (RUP2), AT5G63860 (UVR8).

## Supplemental Data

**Supplemental Figure S1.**
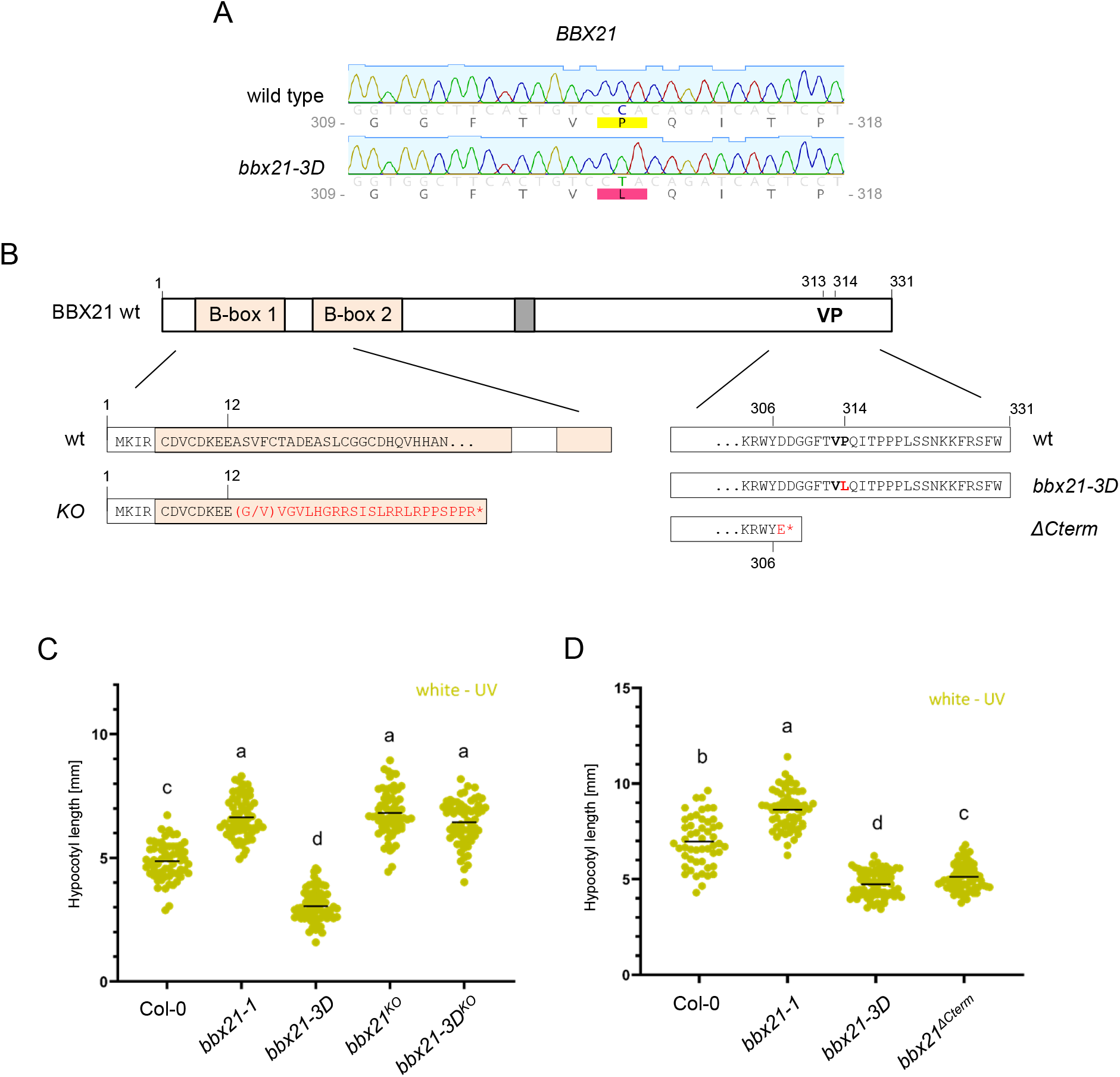
The *bbx21-3D* phenotype is dependent on functional *BBX21*, and can be recapitulated by deleting the C-terminal 25 amino acids that include the VP motif. A, Sequencing chromatograms depicting the C-to-T mutation in *bbx21-3D*, generating the BBX21^P314L^ variant. B, Schematic view of the *BBX21* coding sequence in wild type (wt), *bbx21-3D*, and CRISPR/Cas9-mediated *bbx21* mutant lines (*KO* and *Cterm*). The positions of the two B-box domains, the transactivation domain (grey box), and VP motif are indicated. The altered amino acid residues generated due to the frameshift mutations are indicated in red, with ^*^ representing premature stop codons. KO, knock-out. C, Quantification of hypocotyl lengths of wild type (Col-0), *bbx21-1, bbx21-3D*, and *bbx21* CRISPR/Cas9-mediated knockout lines in the Col-0 (*bbx21*^*KO*^) and *bbx21-3D* (*bbx21-3D*^*KO*^) backgrounds. D, Quantification of hypocotyl lengths of Col-0, *bbx21-1, bbx21-3D*, and a C-terminally truncated CRISPR/Cas9-generated mutant line (*bbx21*^*ΔCterm*^). (C,D) Seedlings were grown for 4 d in white light (values of independent measurements and means as horizontal lines are shown; C, *n* > 60; D, *n* > 50). Shared letters indicate no statistically significant difference between the means (P > 0.05).

**Supplemental Figure S2.**
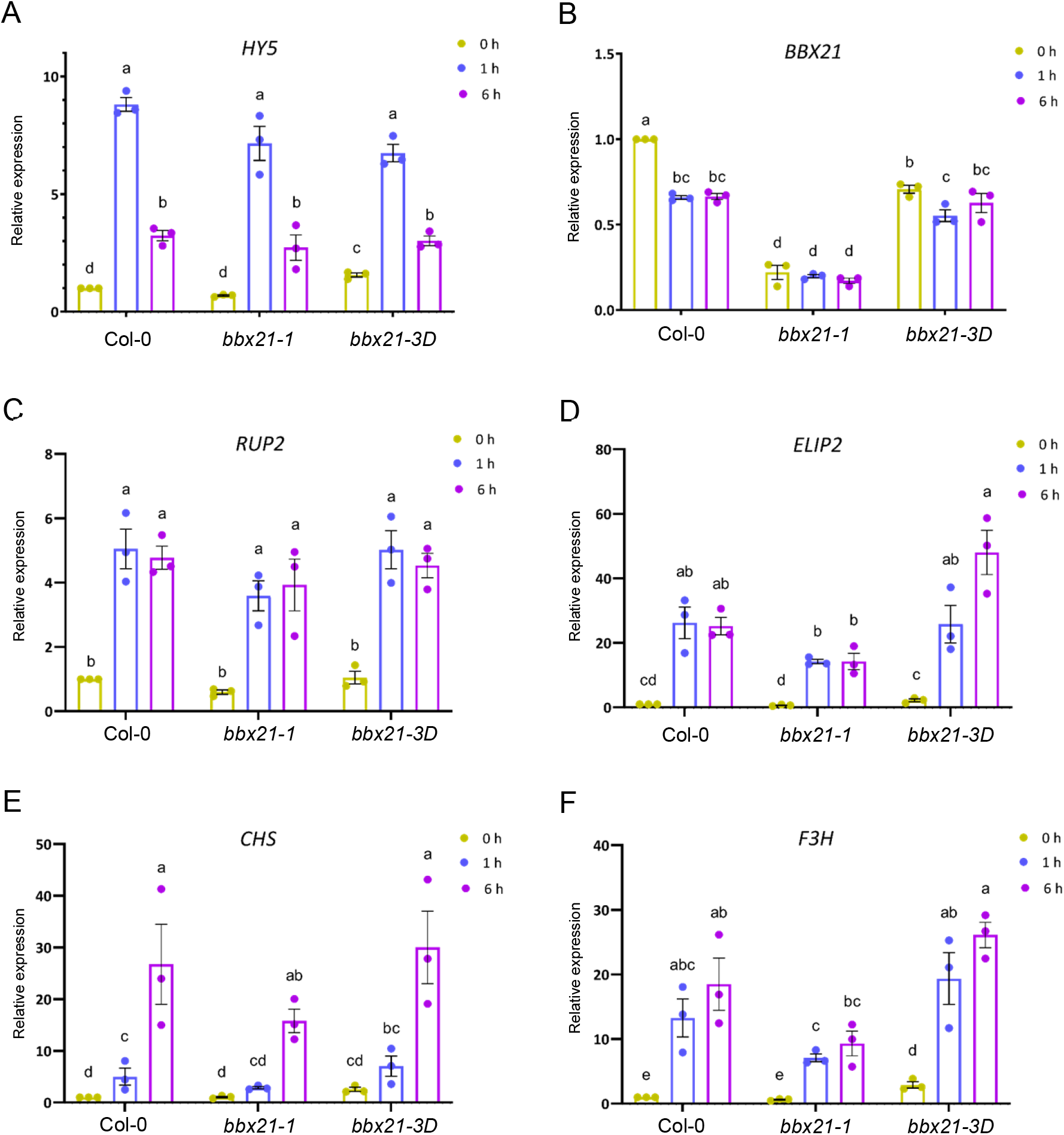
BBX21 promotes the expression of some marker genes in response to UV-B. A–F, RT-qPCR analysis of *HY5* (A), *BBX21* (B), *RUP2* (C), *ELIP2* (D), *CHS* (E), and *F3H* (F) expression in 4-d-old wild-type (Col-0), *bbx21-1*, and *bbx21-3D* seedlings grown in white light and exposed to 1 h and 6 h of supplemental UV-B, or not (0 h). Values of independent measurements, means, and SEM are shown (*n* = 3); shared letters indicate no statistically significant difference between the means (P > 0.05).

**Supplemental Figure S3.**
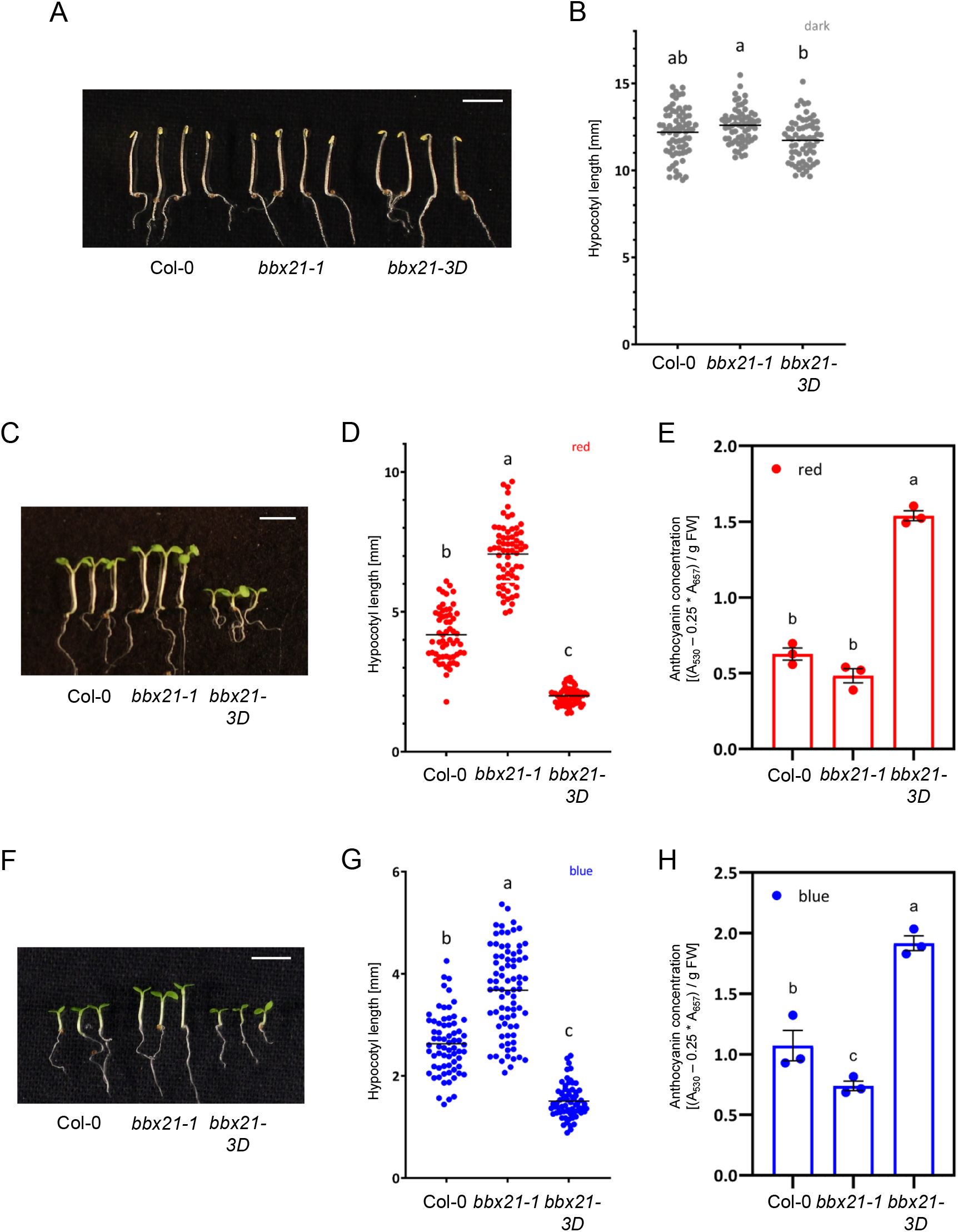
*bbx21-3D* phenotype in darkness and monochromatic red and blue light. A–B, Representative images (A) and quantification of hypocotyl lengths (B) in wild type (Col-0) as well as *bbx21-1* null and *bbx21-3D* gain-of-function mutant seedlings grown in the dark. C–E, Representative images (C), quantification of hypocotyl lengths (D), and quantification of anthocyanin concentrations (E) in Col-0 as well as *bbx21-1* null and *bbx21-3D* gain-of-function mutant seedlings grown under 150 µmol m^-2^ s^-1^ of red light (*n* > 60). F–H, Representative images (F), quantification of hypocotyl lengths (G), and quantification of anthocyanin concentrations (H) in Col-0 as well as *bbx21-1* null and *bbx21-3D* gain-of-function mutant seedlings grown under 50 µmol m^-2^ s^-1^ of blue light. (A,C,F) Scale bar indicates 5 mm. (B,D,G) Values of independent measurements and means as horizontal lines are shown (*n* > 60). (E,H) Values of independent measurements, means, and SEM are shown (*n* = 3). Shared letters indicate no statistically significant difference between the means (P > 0.05).

**Supplemental Figure S4.**
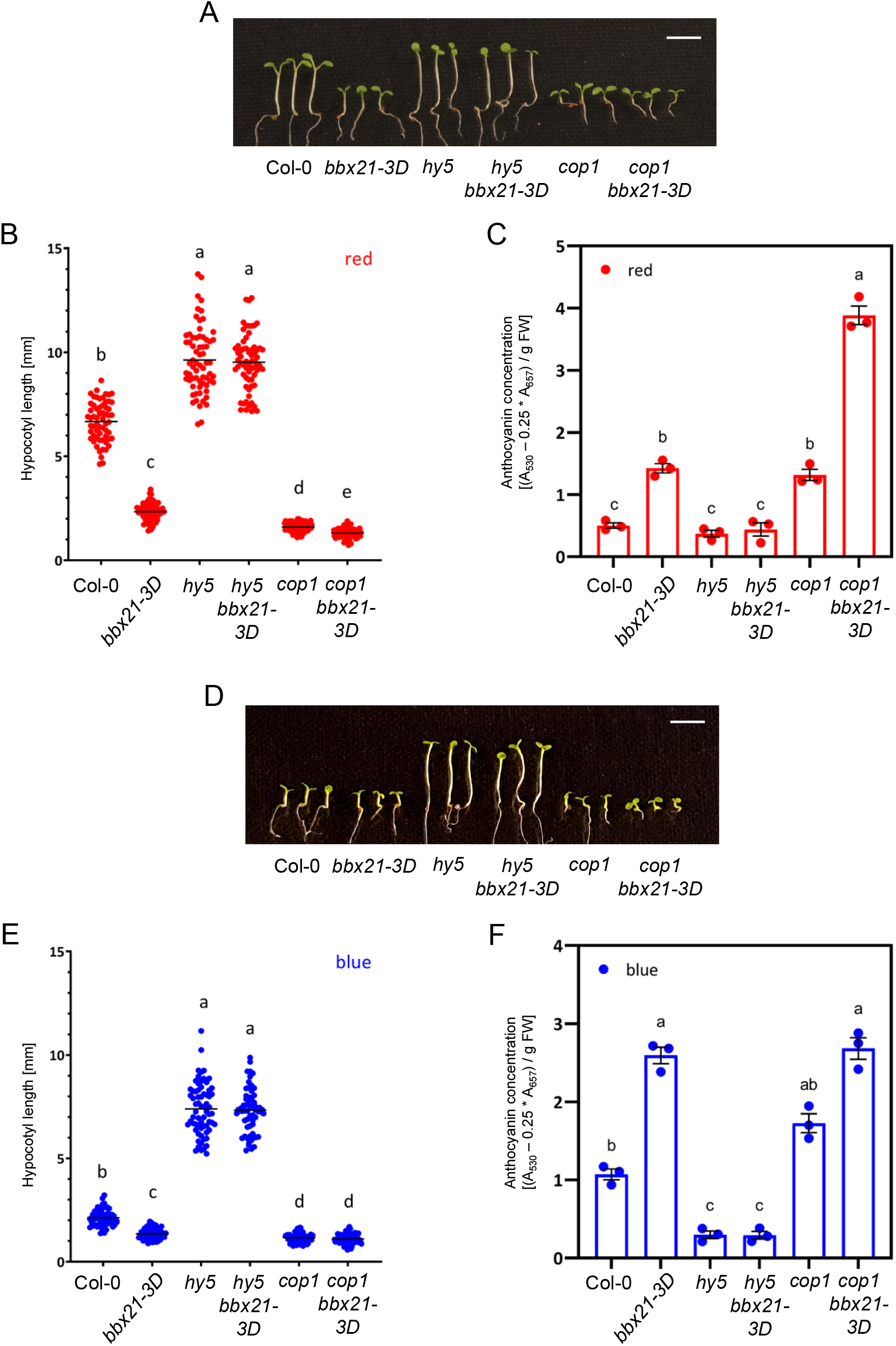
Genetic relationship between *bbx21-3D* and *cop1* and *hy5* in monochromatic red and blue light conditions. A–F, Representative images (A,D; scale bars indicate 5 mm), quantification of hypocotyl lengths (B,E; values of independent measurements and means as horizontal lines are shown; *n* > 60), and quantification of anthocyanin concentrations (C,F; values of independent measurements, means, and SEM are shown; *n* = 3) in wild-type (Col-0), *bbx21-3D, hy5, hy5 bbx21-3D, cop1*, and *cop1 bbx21-3D* seedlings grown under 150 µmol m^-2^ s-^1^ of red light (A–C) or 50 µmol m^-2^ s^-1^ of blue light (D–F). (B,C,E,F) Shared letters indicate no statistically significant difference between the means (P > 0.05).

**Supplemental Figure S5.**
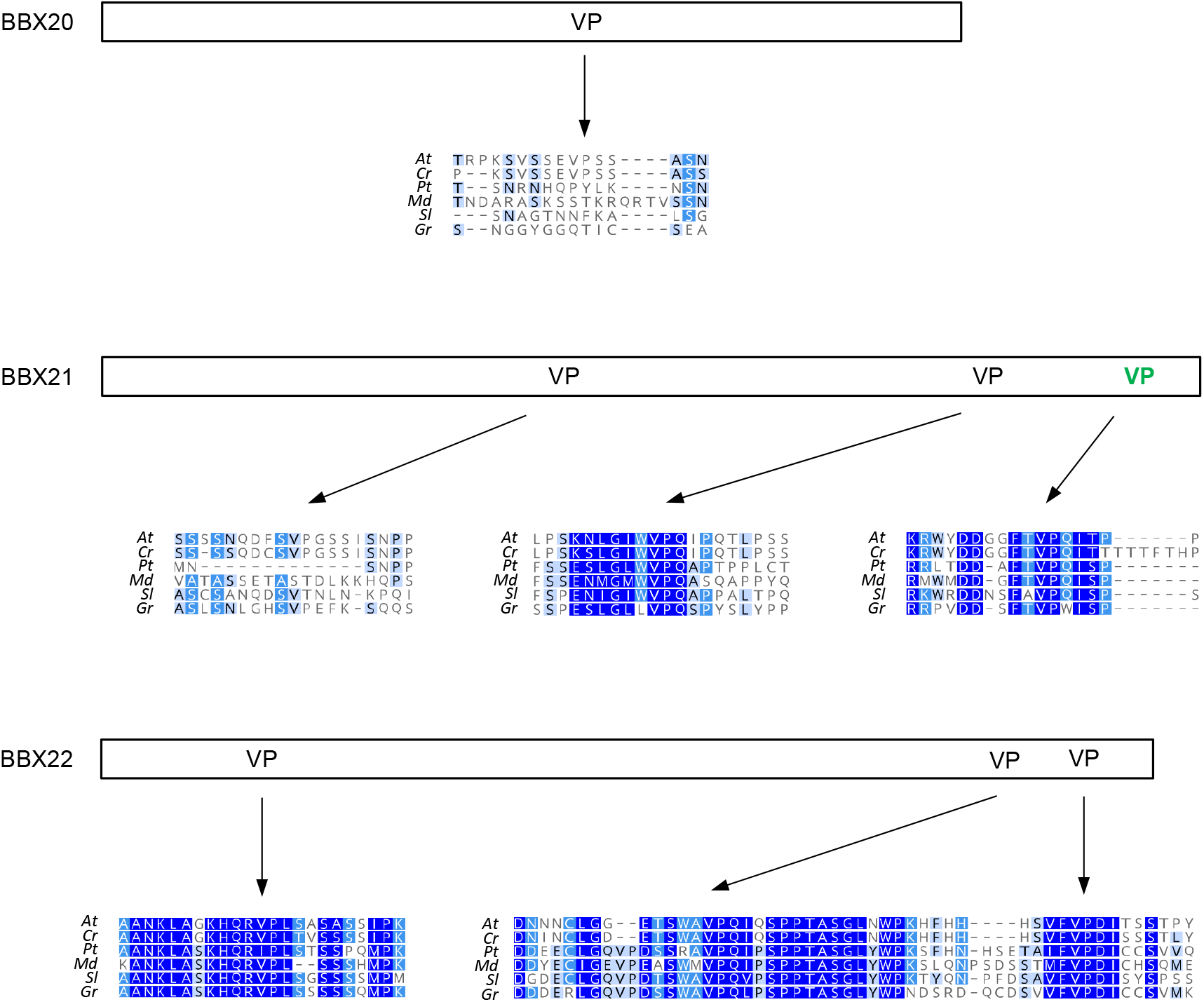
Conservation of putative VP motifs in BBX20, BBX21, and BBX22. Schematic representation of the positions of all VP amino acid pairs in Arabidopsis BBX20, BBX21, and BBX22. In green, reported VP motif in BBX21 (Bursch et al., 2020). For each VP pair, an alignment of homologs, as determined by reciprocal BLASTs on the Phytozome resource (phytozome.jgi.doe.gov) is shown with conservation indicated in blue. Species analysed: *At, Arabidopsis thaliana*; *Cr, Capsella rubella*; *Pt, Populus trichocarpa*; *Md, Malus domestica*; *Sl, Solanum lycopersicum, Gr, Gossypium raimondii*.

**Supplemental Figure S6.**
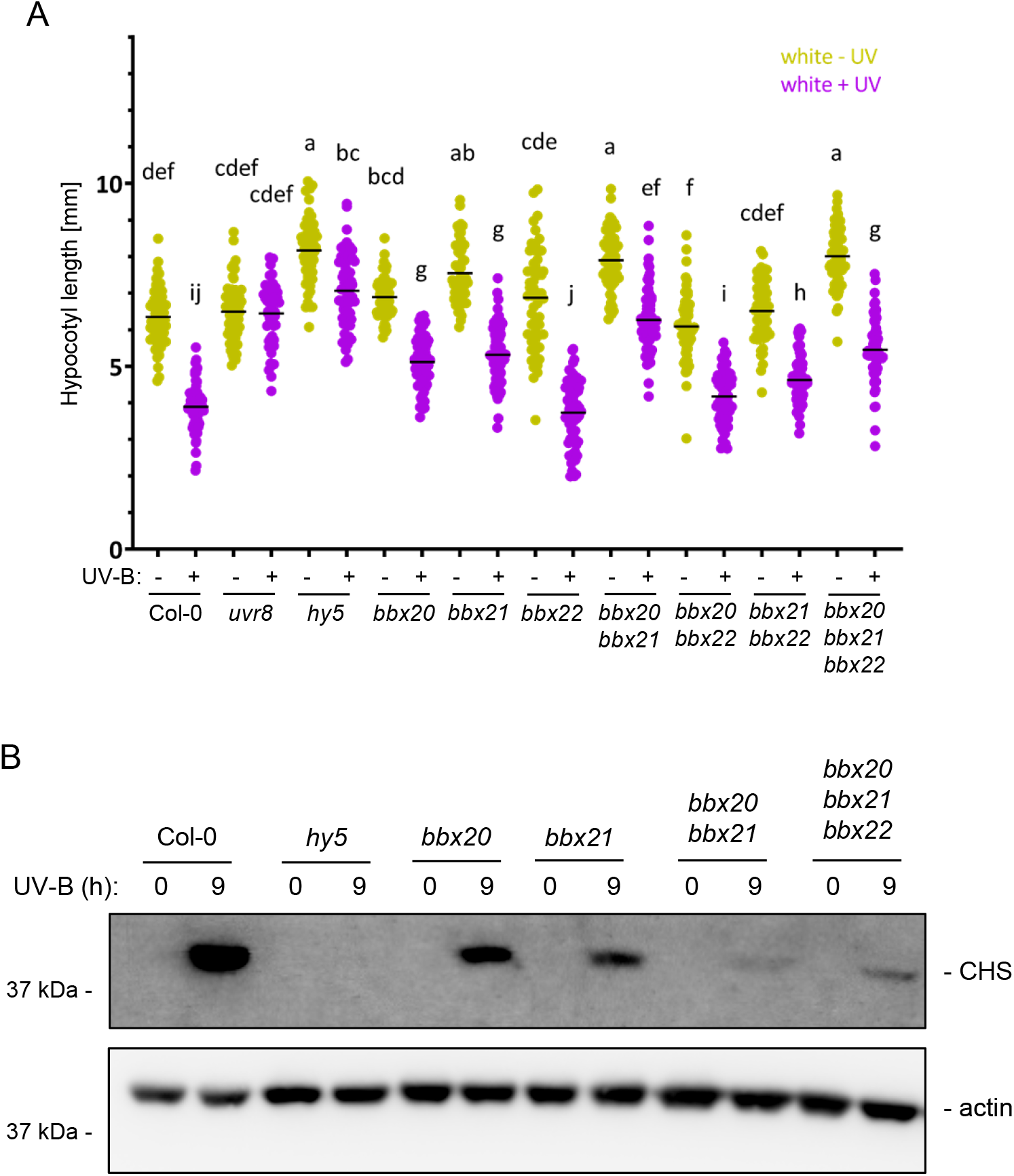
Genetic redundancy between BBX20, BBX21, and BBX22. A, Quantification of hypocotyl lengths of seedlings of wild-type (Col-0), *uvr8-12* (*uvr8*), *hy5-215* (*hy5*), and single and combinatorial mutants of *bbx20, bbx21*, and *bbx22*. Values of independent measurements and means as horizontal lines are shown (*n* > 60). Shared letters indicate no statistically significant difference between the means (P > 0.05). B, Immunoblot analysis of CHS and actin (loading control) levels in Col-0, *hy5*, and single and combinatorial mutants of *bbx20, bbx21*, and *bbx22*. Seedlings were grown for 4 d and treated for 9 h with supplemental UV-B, or not (0).

**Supplemental Table S1.**
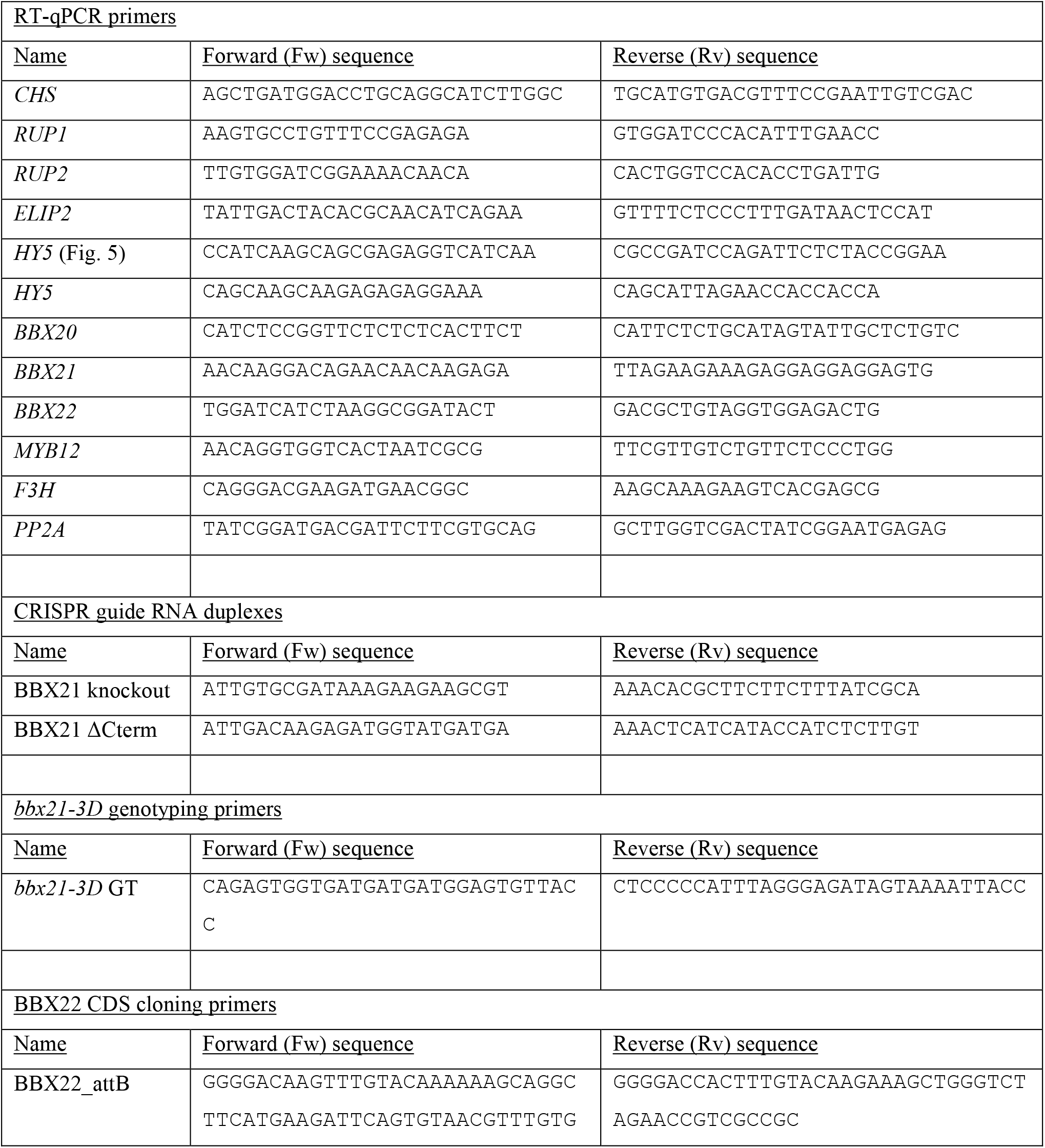
Primers used in this study

## ACKNOWLEDGEMENTS

We would like to thank Emilie Demarsy and Michel Goldschmidt-Clermont for critically reading the manuscript. This work was supported by the University of Geneva, the Swiss National Science Foundation (grant 31003A_175774 to R.U.), and an iGE3 PhD salary award (to R.P.). H.J. was supported by the German Research Foundation (project number 320656366).

## AUTHOR CONTRIBUTIONS

R.P. and R.U. designed the research; T.B.W., R.P., and M.L. performed the experiments; H.J. contributed new tools; T.B.W., R.P., and R.U. analyzed the data and wrote the paper. All authors reviewed and approved the submitted manuscript.

